# Critical Role of P-Glycoprotein-9 in Ivermectin Tolerance in Nematodes

**DOI:** 10.1101/2025.07.10.664073

**Authors:** Clara Blancfuney, Eva Guchen, Marie Garcia, Jean-François Sutra, Felipe Ramon-Portugal, Elise Courtot, Marlène Z Lacroix, Roger Prichard, Anne Lespine, Mélanie Alberich

**Author notes:** Corresponding authors: Mélanie Alberich; Anne Lespine.

## Abstract

Helminth infections in grazing ruminants are of major concern for animal welfare and cause substantial economic losses, prompting the widespread use of ivermectin (IVM). The emergence of IVM resistance, driven by complex and poorly understood mechanisms, increasingly compromises treatment efficacy. Drug efflux transporters, particularly P-glycoproteins (PGPs), are suspected to contribute to resistance. Yet, the study of their individual and functional role is hindered by their diversity in nematodes. This study aimed to dissect the role of specific PGPs in mediating IVM resistance. Thus, *Caenorhabditis elegans* strain IVR10, selected for IVM resistance and reported to overexpress *pgp*s, was used as a model. We generated different IVR10 strains each lacking one of six key *pgp*s, and assessed changes in IVM tolerance. Remarkably, only the deletion of *pgp-9* significantly increased IVM susceptibility. Furthermore, transgenic expression of *Haemonchus contortus pgp-9.1* rescued the resistant phenotype, demonstrating a conserved function across species. To explore drug dynamics, we developed a fluorescent IVM analog, which revealed reduced drug accumulation in IVR10, a phenotype reversed by *pgp-9* deletion. Altogether, these findings show that nematode PGP-9 modulates IVM tolerance by controlling drug efflux and highlight it as a potential therapeutic target.

## Introduction

Helminth infections remain one of the leading causes of morbidity in humans[1,2] and animals. In small ruminants, these infections are of major concern for animal welfare and production[3], and can lead to massive economic losses [4].

Treatments essentially rely on chemicals such as ivermectin (IVM), a broad-spectrum anthelmintic (AH) belonging to the macrocyclic lactone (ML) family. IVM has been massively administered since its market release in 1981 [5] in animals and in humans. This intensive use has inevitably led to the emergence of drug resistance, now undermining control of animal gastro-intestinal parasites, *e.g.*, *Haemonchus contortus* [6]. Moreover, the emergence of resistance in *Onchocerca volvulus* [7], a human filariae helminth treated with IVM, is of growing concern. Considering the lack of development of new pharmaceuticals and the increasing spread of IVM resistance, uncovering its resistance mechanisms stands as an important challenge [8].

IVM resistance seems to be multifactorial, involving overlapping and interacting mechanisms which are poorly understood. Accumulating evidence suggests that the regulation of detoxification systems plays a central role in the development and persistence of resistance in nematodes [9]. A number of studies demonstrated an up-regulation of genes encoding P- Glycoproteins (PGPs), ATP-Binding Cassette efflux pumps of the plasma membrane, in IVM resistant parasites such as *H. contortus* [10–13], *Parascaris univalens* [14] and *Teladorsagia circumincta* [15]. Their expected contribution points to IVM efflux and a protective role against drug accumulation in the nematodes [16]. Among the evidence supporting this role, are the activity modulation of heterologously expressed parasite PGPs in response to AH stimuli [17–20], and their protective role in various toxicity assays [14,21,22]. Furthermore, mammalian PGP inhibitors have been shown to enhance IVM sensitivity in parasitic isolates [23,24]. These data support the notion that helminths PGPs can interact with and transport IVM. However, given the broad repertoire of *pgp* genes in parasites, their individual contributions in this context are still not fully understood.

Studying MLs resistance in parasites proves to be a challenging task because of their complex life cycle and dependence on host propagation. Thus, the free-living nematode *Caenorhabditis elegans* stands out as a powerful well-known model. The development of IVM-resistant strains [25], knock-down and knock-out strains [26] and transcriptional studies [27,28] have been pivotal in understanding of AH action and resistance, and in screening for genes associated with resistance.

In addition to PGPs, other mechanisms may contribute to IVM resistance, notably those involving amphids, chemosensory organs of the nematode. Both IVM-resistant *C. elegans* and *H. contortus* exhibit a dye-filling defect of amphids, suggesting impaired sensory function [27,29,30]. Recent studies have also underscored the importance of NHR-8, a conserved nuclear hormone receptor in *C. elegans* and *H. contortus*, supporting a critical regulatory role of NHR-8 on xenobiotic response pathways [27,31]. Interestingly, *osm-3* mutants display NHR-8 mediated *pgp* overexpression and are drug-resistant [32]. This suggests that PGPs, under the control of NHR-8, coordinate environmental sensing with the regulation of drug resistance mechanisms.

Studies investigating *pgp*s in IVM resistance remain limited to transcriptomics and heterologous expression in models, and their contribution is poorly characterized in IVM- resistant strains. To address this gap, we investigate the role of specific PGPs by selectively targeting genes that are strongly upregulated in the IVM-resistant strain IVR10 [25,27], and downregulated in the *nhr-8* loss-of-function mutant [31], namely: *pgp-1*, *pgp-3*, *pgp-6*, *pgp-9*, *pgp-11* and *pgp-13*. We assessed the impact of individual *pgp* deletions on IVM sensitivity in the IVR10 background. We identified PGP-9 as a key candidate, and further characterized the function of the *H. contortus* ortholog in transgenic worms. Finally, we developed a fluorescent IVM derivative to correlate drug accumulation with the resistant phenotype. This study provides the first direct *in vivo* evidence that PGP-9 from both *C. elegans* and the parasitic nematode *H. contortus* can mediate IVM resistance, establishing its central role in drug efflux and linking PGP activity to reduced drug accumulation at the target organism level.

## Results

### 1. Dominant role of *pgp-9* in IVM tolerance

To evaluate the contribution of each PGPs to IVM tolerance in *C. elegans*, we assessed the IVM sensitivity in IVR10 single knock-out strains for *pgp-1*, *-3*, *-6*, *-9*, *-11*, and *-13* using a larval development assay (LDA). These *pgps* were selected among the fourteen expressed in *C. elegans* based on their overexpression in the IVR10 strain [27] and putative regulation by NHR-8 [31]. IVR10 and N2B were used as resistant and susceptible controls, respectively. Dose-response curves analysis revealed that only the deletions of *pgp-3* and *-9* significantly increased IVM sensitivity in IVR10 (Figure 1A-B). Notably, *pgp-9* deletion resulted in a strong shift to the left of the dose-response curve (Figure 1A), supported by a three-fold reduction in IC_50_ compared to IVR10 (3.98 ± 1.22 *versus* 11.22 ± 1.98 nM, p<0.0001) (Table 1); whereas the effect of *pgp-3* was relatively moderate (6.37 ± 2.71 nM, p<0.001) (Figure 1B, Table 1). This was supported by RFs of 2.5 and 4.0, respectively, underscoring that *pgp-9* deletion had the most substantial, yet partial, effect in reducing IVM-resistance in IVR10. In contrast, the individual knock-out strains for *pgp-1*, *pgp-6*, *pgp-11,* and *pgp-13* did not impact the IVM susceptibility of IVR10, as indicated by IC_50_ values and RFs comparable to those of the resistant parental strain (Table 1).

**Figure 1.**
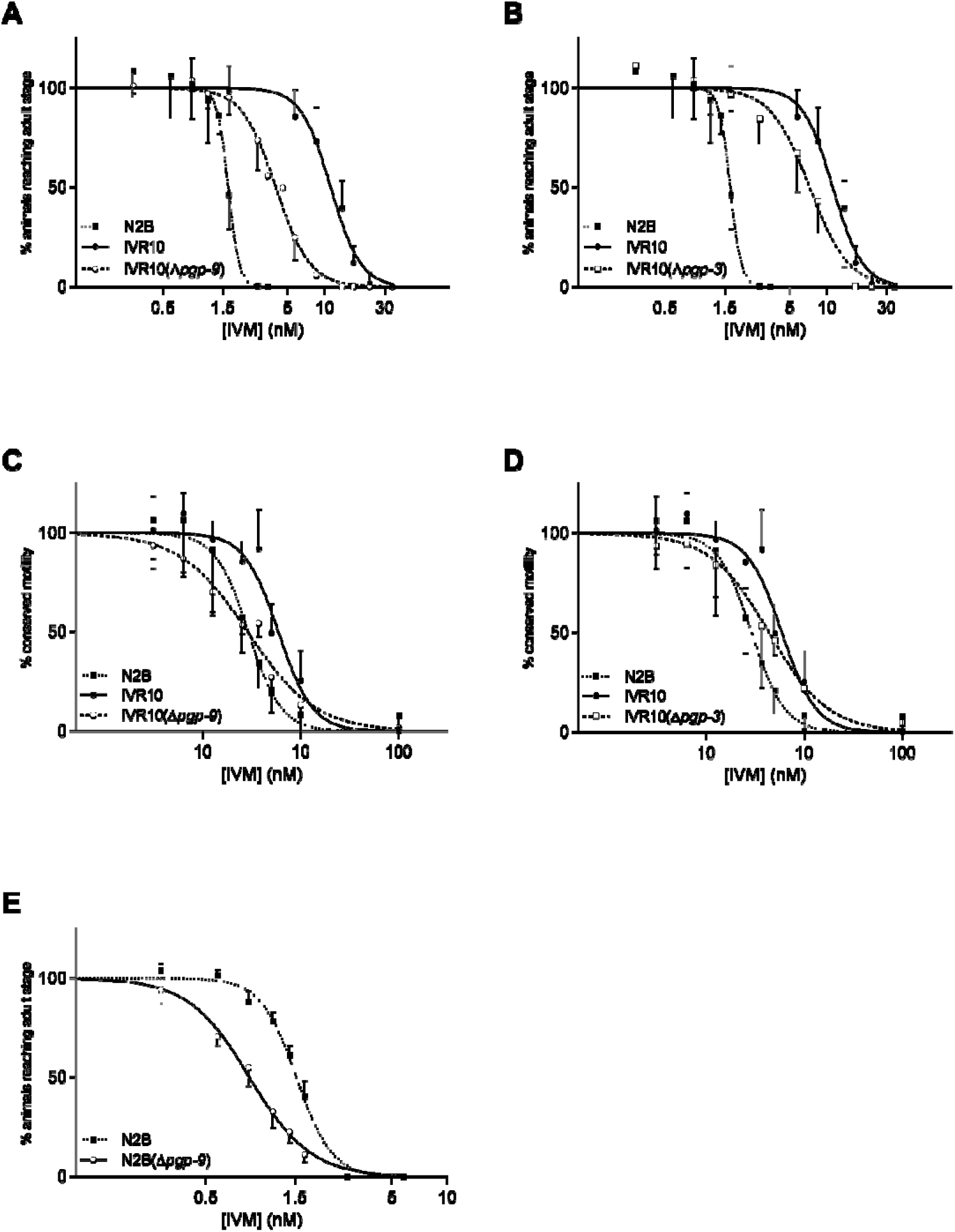
*pgp-9* mediates IVM tolerance. Effects of loss of *pgp-9* and *pgp-3* on tolerance of IVR10 to ivermectin (IVM) in a larval development assay (LDA) (**A** and **B** respectively) and in a motility assay (MA) (**C** and **D** respectively). Effect of loss of *pgp-9* on tolerance of N2B to IVM in a LDA (**E**). Dose-response curves to IVM in LDA and MA are compared to N2B and IVR10 as controls. Values represent the percentage of L1 reaching the young adult stage (LDA) or the percentage of young adults maintaining motility (MA) within the presence of increasing concentrations of IVM. Data are mean ± S.D. from 3-13 independent experiments. IC_50_s for each strain are presented in Table 1, 2 and 3.

**Table 1.** Susceptibilities to ivermectin (IVM) of wild-type Bristol N2 strain, IVM selected strain (IVR10), and genetically modified IVR10 for which *pgp-1*, *-3*, *-6*, *-9*, *-11,* and *-13* have been deleted on larval development assay (LDA). IC_50_: inhibitory concentration 50%. RF: resistance factor, fold resistance relative to the N2B.

| Strain | Mean IC<sub>50</sub> (nM) | RF |
| --- | --- | --- |
|  | ± S.D. (no. of experiments) |  |
| Bristol N2 | $1.61 \pm 0.15$ (3) | - |
| IVR10 | 11.22 ± 1.98 (13) | 6.7 |
| IVR10($\Delta pgp-1$) | 9.20 ± 0.72 (3) | 5.7 |
| IVR10($\Delta pgp-3$) | 6.37 ± 2.71 (4)<sup>b</sup> | 4.0 |
| IVR10($\Delta pgp-6$) | 9.43 ± 2.70 (4) | 5.9 |
| IVR10($\Delta pgp-9$) | 3.98 ± 1.22 (4)<sup>a</sup> | 2.5 |
| IVR10($\Delta pgp-11$) | 8.93 ± 1.92 (4) | 5.6 |
| IVR10($\Delta pgp-13$) | 9.97 ± 1.55 (4) | 6.2 |

<sup>a</sup> p< 0.0001; <sup>b</sup> p< 0.001 knock-out strain *versus* IVR10 (one-way ANOVA followed Dunnett's multiple comparisons test).

Since IVM targets glutamate-gated chloride channels (GluCls) [33] and induces paralysis in nematodes, we conducted a motility assay (MA) to compare worm motion in the different strains. As expected, N2B and IVR10 exhibited markedly different motility responses to IVM with IC_50_ values of 28.50 ± 7.51 and 61.88 ± 16.83 nM, respectively (p<0.01), corresponding to a 2.2 RF (Figure 1, C-D, Table 2). Consistent with the LDA results, *pgp-9* deletion led to a pronounced increase in IVM sensitivity, with a two-fold reduction in IC_50_ compared to IVR10 (29.14 ± 4.02 *versus* 61.88 ± 16.83 nM, p<0.05), and a shift to the left of the dose-response curve (Figure 1C, Table 2). Importantly, the RF of IVR10(Δ*pgp-9*) appeared to be equal to 1.0, revealing that *pgp-9* deletion completely restored IVM sensibility. Meanwhile, deletion of *pgp-3* in IVR10 only led to a modest shift in IVM sensitivity (41.34 ± 12.59 *versus* 61.88 ± 16.83 nM, ns) (Figure 1D, Table 2). Finally, we confirmed that deletions of *pgp-1*, *pgp-6*, *pgp-11,* and *pgp-13* did not impact IVM sensitivity in IVR10, as their deletion did not alter either the motility phenotype, as reflected by their respective RFs comparable to that of the resistant IVR10 strains (Table 2).

**Table 2.**
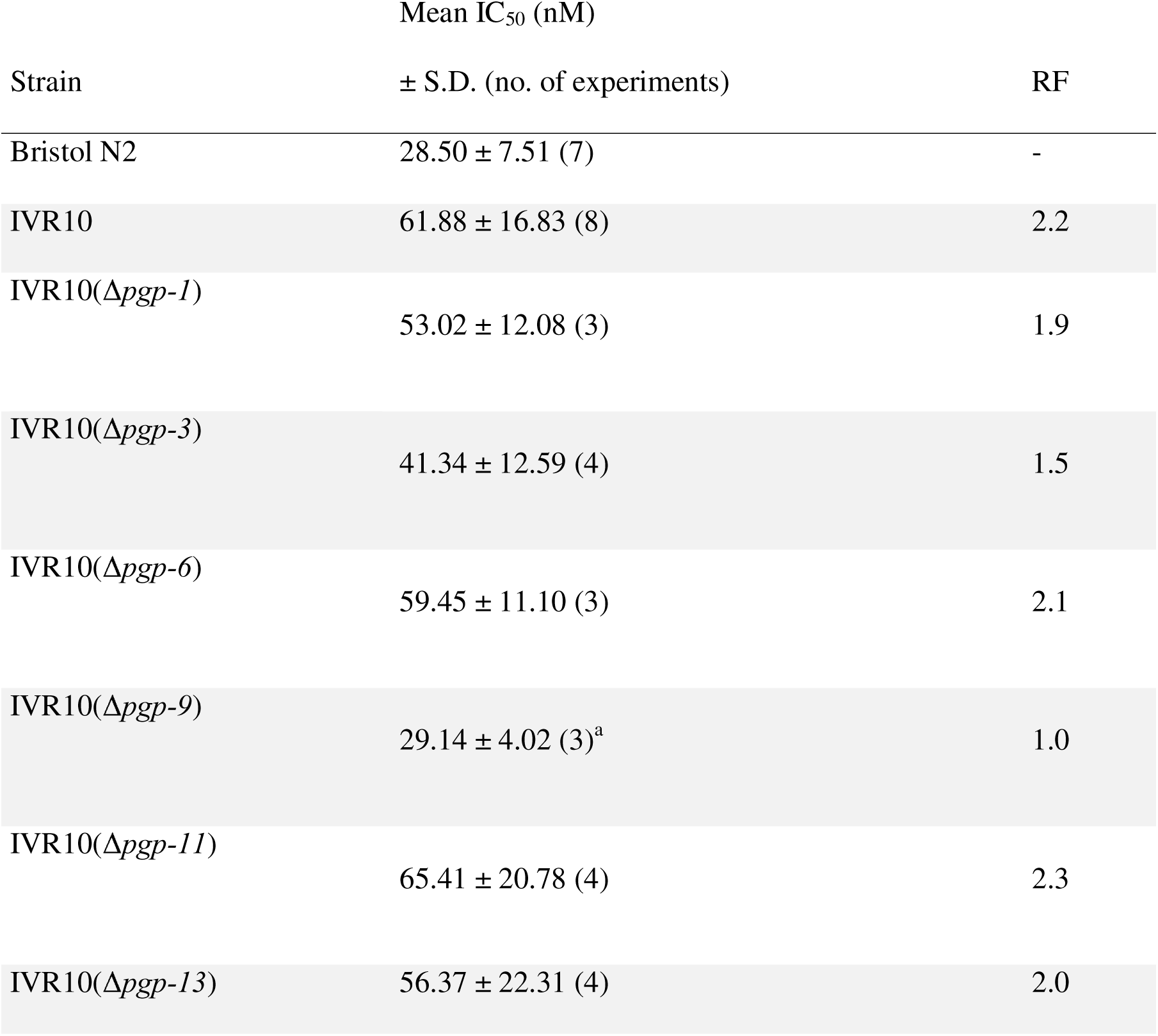
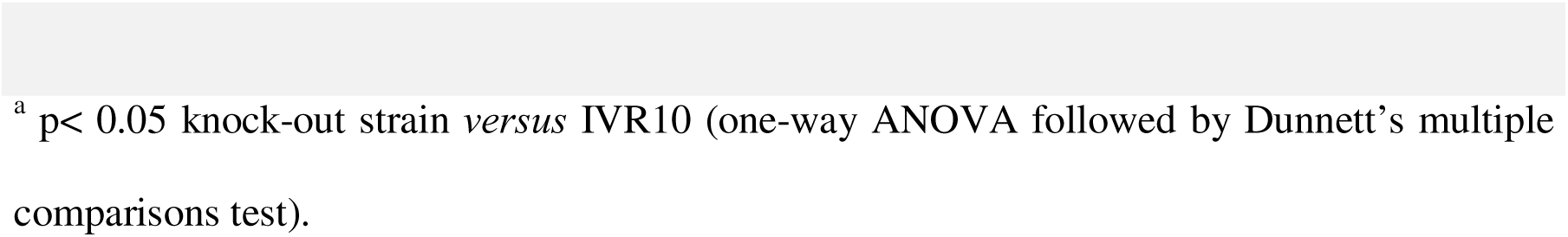
Susceptibilities to ivermectin (IVM) of wild-type Bristol N2 strain, IVM selected (IVR10) strains, and genetically modified IVR10 for which *pgp-1*, *-3*, *-6*, *-9*, *-11,* and *-13* have been deleted on motility assay (MA). IC_50_: inhibitory concentration 50%. RF: resistance factor.

| Strain | Mean IC<sub>50</sub> (nM) | RF |
| --- | --- | --- |
|  | ± S.D. (no. of experiments) |  |
| Bristol N2 | $28.50 \pm 7.51$ (7) | - |
| IVR10 | $61.88 \pm 16.83$ (8) | 2.2 |
| IVR10($\Delta pgp-1$) | $53.02 \pm 12.08$ (3) | 1.9 |
| IVR10($\Delta pgp-3$) | $41.34 \pm 12.59$ (4) | 1.5 |
| IVR10($\Delta pgp-6$) | $59.45 \pm 11.10$ (3) | 2.1 |
| IVR10($\Delta pgp-9$) | $29.14 \pm 4.02$ (3)<sup>a</sup> | 1.0 |
| IVR10($\Delta pgp-11$) | $65.41 \pm 20.78$ (4) | 2.3 |
| IVR10($\Delta pgp-13$) | $56.37 \pm 22.31$ (4) | 2.0 |

<sup>a</sup> $p < 0.05$ knock-out strain *versus* IVR10 (one-way ANOVA followed by Dunnett's multiple comparisons test).

We next extended our analysis to two additional IVM-resistant strains previously and independently generated in our laboratory by step-wise exposure of N2B to increasing IVM concentrations: IVR10-2014 and IVR10-2022. *Pgp-9* was silenced in both strains by feeding with RNAi, and IVM sensitivity was assessed using LDA. We first validated *pgp-9* silencing by confirming a strong decrease (around 80%) in *pgp-9* transcripts using RT-qPCR (S2 Figure). IC_50_ values and dose-response curves were compared to controls fed with HT115- control (expressing the empty RNAi vector). IVR10-2014 and IVR10-2022 exhibited phenotypes similar to the standard strain IVR10 on HT115-control and therefore reflective of worms resistant to IVM, as supported by their respective IC_50_ values (S3 Table). *Pgp-9* silencing induced a moderate but consistent shift to the left of the dose-response curve of each IVM-resistant strain (S1 Figure). This was supported by a significant decrease, up to 1.8-fold, of the IC_50_ values compared to the control RNAi (S1 Figure 1, S3 Table).

Finally, to assess the role of *pgp-9* in modulating IVM tolerance in the susceptible strain N2B, we investigated IVM sensitivity in a mutant strain for *pgp-9*. LDA revealed a significant increase in IVM sensitivity supported by a shift to the left of the dose-response curve (Figure 1 E) and by a 1.8-fold reduction in IC_50_ values for N2B(Δ*pgp-9*) compared to N2B (Table 3) (0.86 ± 0.10 nM *versus* 1.54 ± 0.07, respectively, p< 0.001).

**Table 3.** Susceptibilities to ivermectin (IVM) of wild-type Bristol N2 strain and *pgp-9* loss-of-function N2B in larval development assay (LDA). IC_50_: Inhibitory concentration 50%.

| Strain | Mean IC<sub>50</sub> (nM) ± SD (no. of experiments) |
| --- | --- |
| Bristol N2 | 1.54 ± 0.07 (3) |
| N2B(Δ*pgp-9*) | 0.86 ± 0.10 (4)<sup>a</sup> |

<sup>a</sup> p < 0.001, N2B( $\Delta$ *pgp-9*) versus N2B (parametric unpaired t-test).

Together, these results demonstrate that *pgp-9* plays a central role in maintaining IVM tolerance across multiple independently selected resistant *C. elegans* strains and in the wild-type susceptible strain, and demonstrate that its deletion increases both developmental and motility-based IVM susceptibility.

### 2. Functional conservation of PGP-9 in *H. contortus*

We next investigated whether the function of PGP-9 in IVM resistance is conserved across parasitic species, particularly in the gastrointestinal nematode *H. contortus*. We assessed whether *Hco-pgp-9.1* (also referred to as *Hco-pgp-9*) can compensate for the loss of *Cel-pgp- 9* and restore IVM tolerance in the IVR10(Δ*pgp-9*) *C. elegans* strain.

To better understand the potential functional conservation of these transporters, we first assessed the identity between the two transporters. Given the high conservation of their nucleotide-binding domains (NBDs), which are involved in ATP binding, we excluded these regions from both sequences to focus on the transmembrane domains (TMDs), which are critical for ligand binding and specificity. Partial protein sequence alignment revealed a 63.7% identity between *Cel*-PGP-9 and *Hco*-PGP-9.1 (S3 Figure), supporting a conserved functional role.

We then expressed *Hco-pgp-9.1* in the IVR10 *pgp-9* knock-out strain under *Cel-pgp-9* promoter. Three independent rescue strains were studied: IVR10(Δ*pgp-9*)-R#1, IVR10(Δ*pgp- 9*)-R#2 and IVR10(Δ*pgp-9*)-R#3. Transcription of *Hco-pgp-9.1* transgene was confirmed by single worm RT-qPCR (S4 Figure) in the three independent rescue strains, and compared to *Cel-pgp-9* mRNA expression in IVR10. No statistical differences were observed, suggesting that the transgene expression in transgenic worms was comparable to the physiological expression of *Cel-pgp-9* in their IVR10 background counterpart.

As expected, the LDA demonstrated that worms lacking the *Hco-pgp-9.1* transgene, *i.e.*, IVR10(Δ*pgp-9*), showed low tolerance to IVM. Indeed, at 4 nM, only 45 to 57% of the population reached the adult stage (Figure 2). At 8 nM, worm development was nearly completely suppressed, in line with the previously determined IC_50_ of 3.99 nM for IVR10(Δ*pgp-9*). In contrast, worms expressing the *Hco-pgp-9* transgene consistently exhibited a significantly higher tolerance to the drug, as revealed by a significantly increased percentage of development in the presence of IVM at 4, 8 and 10 nM (Figure 2, p values < 0.01). At 4 nM, the percentage of development was of 99% (IVR10(Δ*pgp-9*)-R#1), 107% (IVR10(Δ*pgp-9*)-R#2) and 90% (IVR10(Δ*pgp-9*)-R#3). At 8 nM, 71%, 81% and 73%, of the rescue IVR10(Δ*pgp-9*)-R#1, IVR10(Δ*pgp-9*)-R#2 and IVR10(Δ*pgp-9*)-R#3, respectively, reached adulthood. Finally, at 10 nM, 67% (IVR10(Δ*pgp-9*)-R#1), 70% (IVR10(Δ*pgp-9*)- R#2) and 59% (IVR10(Δ*pgp-9*)-R#3) were adults and exhibited an IVM-resistant phenotype equivalent to the original IVR10 strain, whose IC_50_ is of 11.22 nM.

**Figure 2.**
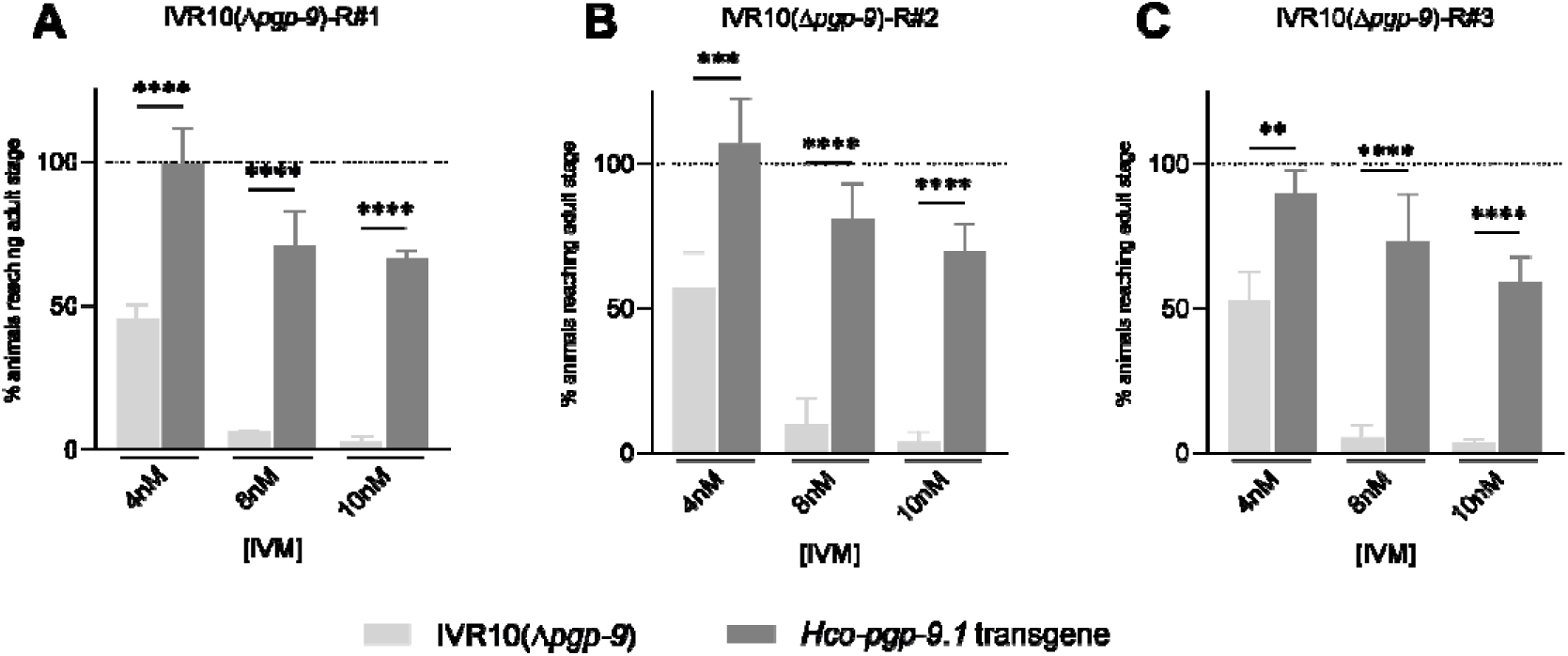
Impact of *Hco-pgp-9.1* rescue in IVR10(Δ*pgp*-9) on IVM tolerance. Effect of transgene expression of *Hco-pgp-9.1* on the tolerance of IVR10(Δ*pgp*-9) to ivermectin (IVM) in a modified larval development assay (LDA). This assay was performed on three independent transgenic strains expressing the *Hco-pgp-9.1* transgene: (**A**) IVR10(Δ*pgp-9*)-R#1, (**B**) IVR10(Δ*pgp-9*)-R#2, and (**C**) IVR10(Δ*pgp-9*)-R#3. For each IVM concentration, the number of adult worms negative (*i.e.*, IVR10(Δ*pgp*-9)) and positive (*i.e.*, *Hco-pgp-9.1* transgene) for the extrachromosomal array was normalized to the number of adult worms of each population present in the DMSO well (full development, dotted line) and expressed as a percentage. The percentages of transgene animals reaching the adult stage for all concentrations were then statistically compared with those of the internal control strain, IVR10(Δ*pgp-9*) (negative for transgene) (two-way anova followed by Sidak’s multiple comparisons test). Data are mean ± S.D. from 3 independent experiments. Transgenic strain genotype: IVR10[*Cel-pgp-9* -; *pCel-pgp-9::Hco-pgp-9.1::SL2::mCh::Cel-unc-54*; pPD118.3].

Overall, these findings demonstrate that the heterologous expression of *Hco-pgp-9* is sufficient to restore a phenotype equivalent to the IVM-resistant one in IVR10(Δ*pgp-9*), indicating functional conservation between *Hco*-PGP-9.1 and *Cel-*PGP-9.

### 3. PGP-9 function is critical in IVM clearance

To further explore the role of *pgp-9* in IVM resistance, we developed a fluorescent analog of IVM, F-IVM. This probe enabled direct visualization and quantification of drug accumulation in *C. elegans*. The synthesis of F-IVM (Figure 3A) yielded a product with 99.9% purity, mainly consisting of two known isomers (B1a: 95.4%, B1b: 2.8%), with minor impurities (1.8%) and traces of native IVM (0.04%). Mass spectrometry comparison of IVM and F-IVM spectra (Figure 3B) demonstrated that the structure of F-IVM closely resembles that of native IVM. The primary difference lies in the loss of two hydroxyl groups from the tetrahydrofuran ring, resulting in the formation of a benzofuran moiety. This structural change enables electron delocalization due to the presence of conjugated bonds, thus conferring fluorescence to the molecule. F-IVM was stable for over 18 hours at room temperature (22 °C) in DMSO (10 mM) and maintained stability for more than five weeks at –20 °C, as well as during long-term storage at –80 °C in DMSO. Efficacy assessment by LDA showed that F-IVM has significantly reduced toxicity compared to native IVM (Figure 3C). The liquid LDA revealed that the susceptible N2B strain exhibited an IC of 2.70 ± 0.70 µM for F-IVM, approximately 1000-fold higher than for IVM. The resistant IVR10 strain demonstrated cross-resistance to F-IVM with an IC of 9.97 ± 0.76 µM (RF of 2.58). We next investigated the impact of F-IVM on the motility of both strains (Figure 3D). Interestingly, F-IVM had no impact on worm motility of both susceptible and IVM-resistant *C. elegans* in the range of investigated concentrations (up to 150 µM) compared to IVM. This fluorescent probe thus offers a powerful tool to investigate drug distribution and accumulation, enabling detailed studies of PGP-mediated IVM resistance mechanisms in *C. elegans*.

**Figure 3.**
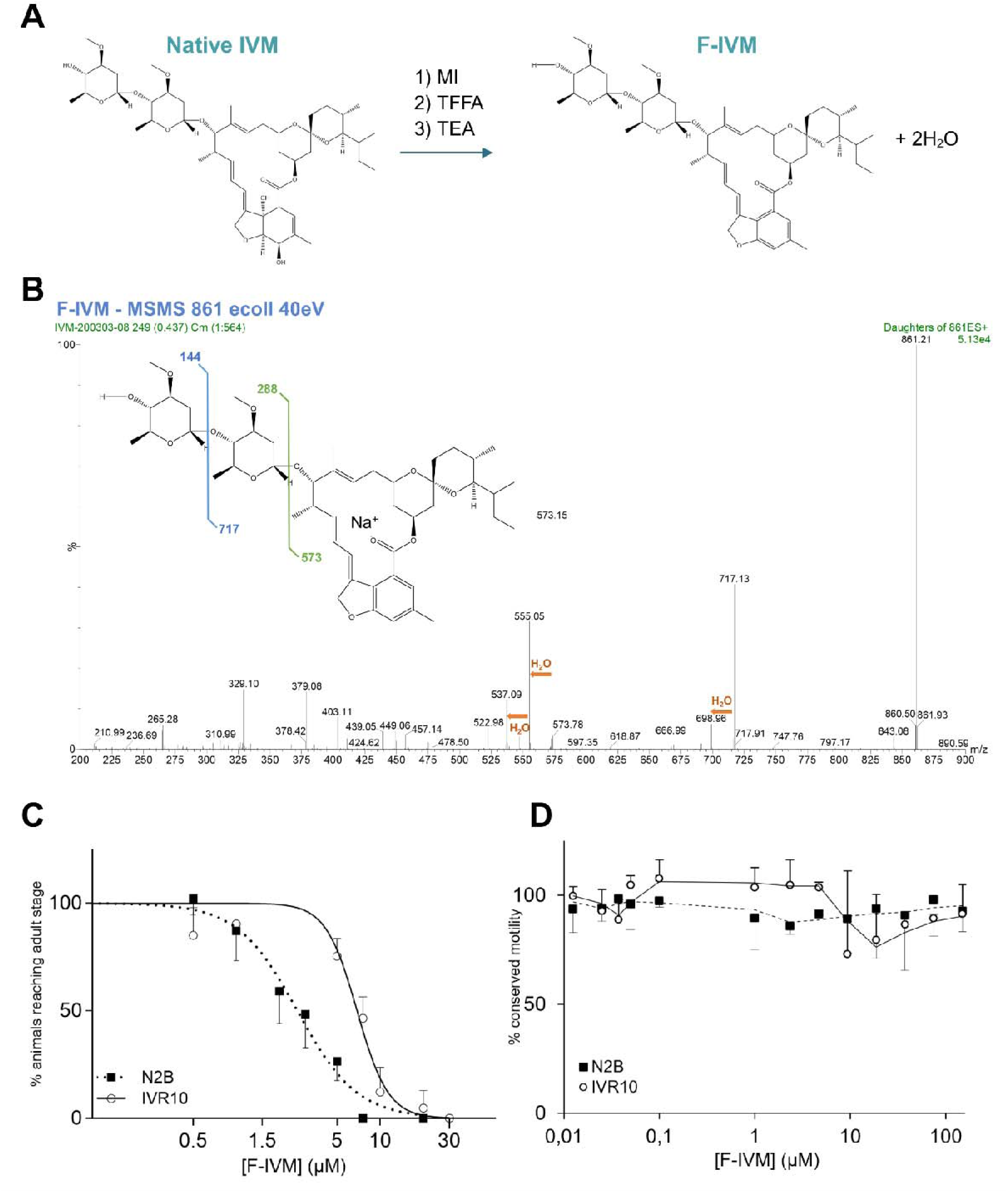
Synthesis, structural characterization, and *in vivo* toxicity assessment of fluorescent ivermectin (F-IVM). (**A**) Schematic representation of the chemical synthesis of F-IVM from native IVM. MI (1-N-methyl-imidazole), TFA (anhydrous trifluoroacetic acid), and TEA (triethylamine). (**B**) Daughter scan mass spectrum of F-IVM showing strong similarity to native IVM, confirming structural conservation. (**C**) Larval development assay (LDA) in *C. elegans* wild-type (N2B) and ivermectin-resistant (IVR10) strains. Dose– response curves show the percentage of L1 larvae reaching the young adult stage at increasing concentrations of F-IVM. (**D**) Motility assay (MA) assessing locomotor activity of young adult worms after 240 min exposure to F-IVM (0.003–150 μM) at 21 °C, using the WMicroTracker system. Data are presented as mean ± S.D. from three independent experiments.

In that context, animals were exposed to the probe to examine PGP-9 function in F-IVM transport comparing IVR10(Δ*pgp-9*) and IVR10, with N2B as a control. All three strains showed consistent accumulation of the fluorescent probe in the pharynx, particularly in the posterior pharyngeal valve (Figure 4A). A closer examination of N2B and IVR10(Δ*pgp-9*) confirmed that the strains displayed a strong signal in the terminal bulbs of the pharynx, as well as the pharyngeal-intestinal valve (Figure 4B), probably leading to diffusion into pharyngeal muscles.

**Figure 4:**
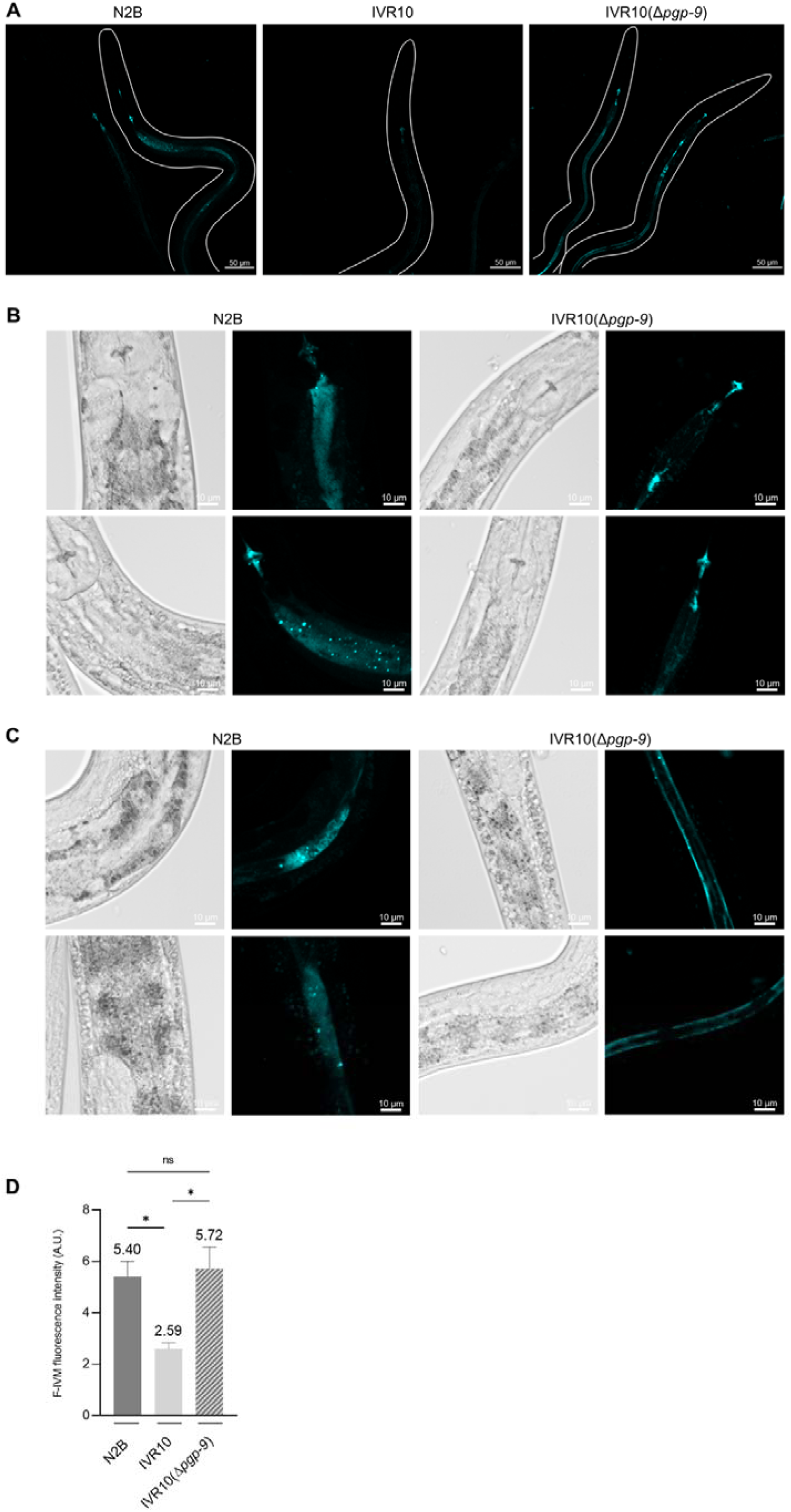
*pgp-9* modulates F-IVM accumulation in IVR10. (**A**) Fluorescent-ivermectin (F-IVM) accumulation in N2 Bristol, IVM-resistant IVR10 and IVR10 *pgp-9* knock-out, IVR10(Δ*pgp-9*). Adult *C. elegans* worms were exposed to F-IVM and then examined by confocal microscopy with a x20 air objective to visualize the fluorescent probe accumulation. Only the fluorescence channel is shown. (**B**) Magnified images of the probe localization around the pharyngeal-intestinal valve in N2B and IVR10(Δ*pgp-9*). (**C**) Magnified images of the probe localization in the intestine in N2B and IVR10(Δ*pgp-9*). Brightfield images are included to aid anatomical interpretation. (**D**) Functional transport assay using IVM fluorescent probe, F-IVM, in N2B, IVR10, and IVR10(Δ*pgp-9*). For each strain, accumulation was quantified in approx. 20-30 adults and experiments were repeated 3 times. Fluorescence was normalized with an untreated population of worms. Data are mean ± S.E.M. * p<0.05 (one-way ANOVA with Tukey’s post-hoc test).

In N2B, F-IVM also accumulated in a punctate pattern at the intestinal region, indicating possible accumulation in intestinal cells or intracellular organelles (Figure 4C), while IVR10 exhibited reduced fluorescence, with minimal or no detectable signal in the intestine (Figure 4A). In contrast, deletion of *pgp-9* in IVR10 restored F-IVM accumulation in the gut, though the distribution pattern differed from that observed in N2B. In IVR10(Δ*pgp-9*), the fluorescent signal was predominantly aligned along the intestinal tract, likely corresponding to accumulation at the apical membrane of intestinal cells (Figure 4C).

Fluorescence intensity measured for each strain supported these observations (Figure 4D). Consistent with our hypothesis, the IVR10 strain exhibited significantly reduced F-IVM accumulation, with a two-fold decrease compared to N2B (2.59 ± 0.25 *versus* 5.40 ± 0.60, p<0.05). Importantly, *pgp-9* deletion in IVR10 restored F-IVM accumulation to levels comparable to the N2B susceptible strain (5.72 ± 0.84 *versus* 5.40 ± 0.60, ns). Taken together, these results show a strong relationship between F-IVM accumulation and the level of IVM resistance in each strain, highlighting PGP-9 as a critical determinant of F-IVM accumulation in the resistant model IVR10.

### 4. Strategic tissue localization of PGP-9

To further elucidate the physiological contexts in which *pgp-9* deletion leads to increased F- IVM accumulation, we examined the tissue-specific expression pattern of the *pgp-9* gene in N2B(Δ*pgp-9*) and IVR10(*pgp-9*) using a *pgp-9* promoter-mCherry (mCh) fusion construct. mCh was localized in the pharyngeal bulbs (anterior and posterior), as well as in the intestine throughout all life stages (L1 to adulthood) (Figure 5). Interestingly, mCh expression was also detected at the periphery of the pharynx muscle, in an area where neuron cells, and notably amphids, are typically localized (“N” arrow in Figure 5). Expression patterns were similar in N2B (Figure 5A) and IVR10 (Figure 5B), underlining a conserved tissue-specific localization between these two strains. In summary, our data suggest that PGP-9 is mainly expressed in the pharynx, in the gut, and neurons, independently of the resistance status.

**Figure 5.**
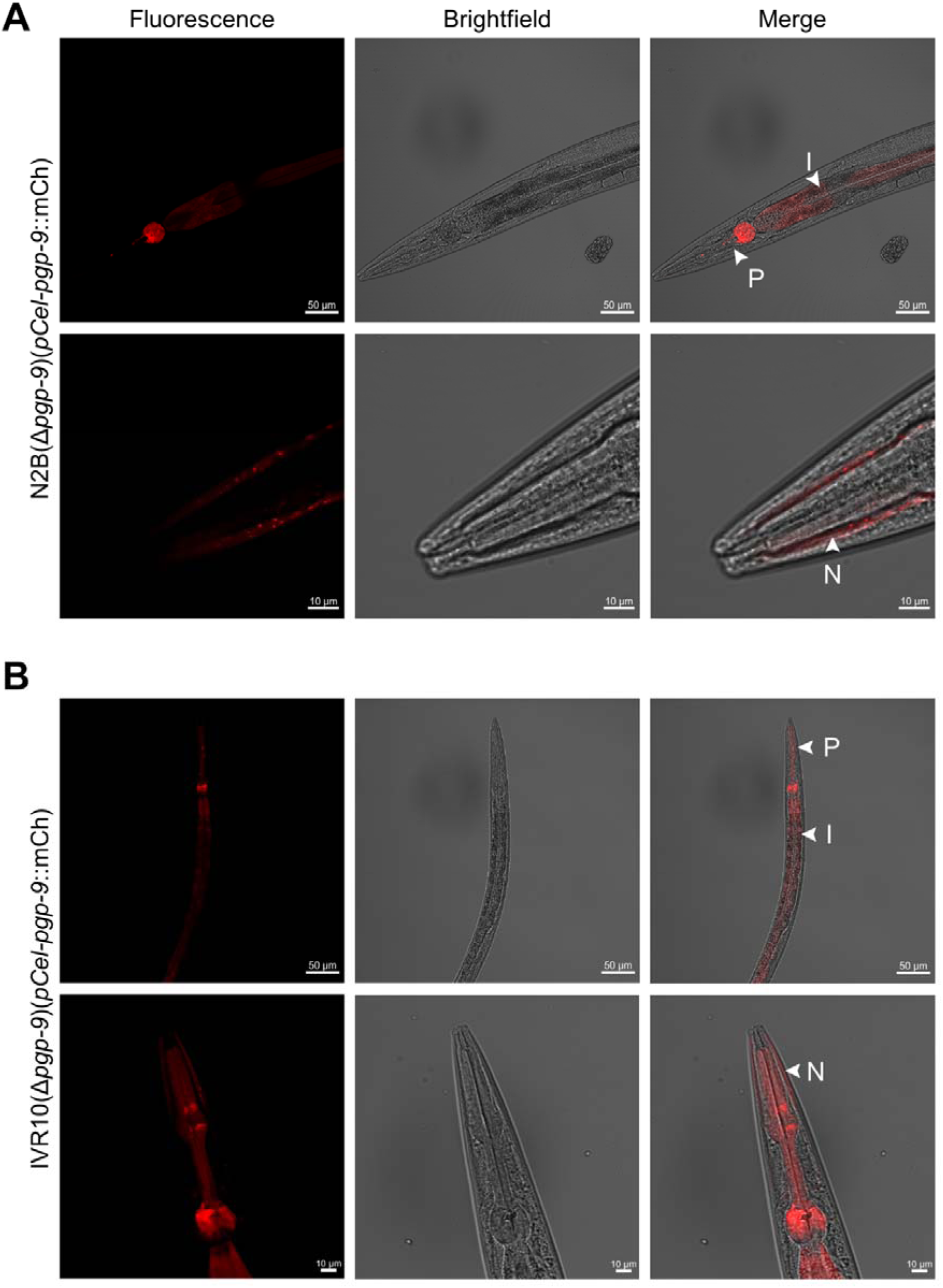
mCh driven expression by *pgp-9* promoter. Expression patterns were observed in (**A**) N2B(Δ*pgp-9*) and (**B**) IVR10(Δ*pgp-9*). Strains were observed using confocal fluorescence microscopy. Expression was observed in all life stages, mainly in the pharynx anterior and posterior bulbs and in the intestine, as shown by the arrows. Strains also demonstrated expression of mCh in the area where neuron cells are localized. Arrows indicate: P: pharynx; I: intestine; N: neurons. Transgenic strain genotype: N2B[*Cel-pgp-9* -; *pCel-pgp-9::SL2::mCh::Cel-unc-54*; pSC391]; IVR10[*Cel-pgp-9* -; *pCel-pgp-9::SL2::mCh::Cel-unc-54*; pPD118.3].

## Discussion

Understanding the functional basis of IVM resistance in nematodes remains a major challenge. In this study, we provide new insights into the role of nematode PGPs in this context. We relied on the *C. elegans* IVR10 strain, generated by IVM stepwise selection pressure, approaching real-life patterns of drug exposure and offering a therapeutically relevant resistant model. Our strategy involved targeted deletions of *pgp* genes in *C. elegans* directly within the IVM-resistant background using CRISPR-Cas9. Thereby, we generated IVR10-derived strains with full-length individual deletions of *pgp-1, pgp-3, pgp-6, pgp-9, pgp-11,* and *pgp-13*. These genes were previously reported to be upregulated in IVR10 [27] and down-regulated in the *nhr-8* mutant [31], making them prime targets of interest in the context of IVM resistance. This approach allowed us to directly assess the contribution of each transporter to the IVM-resistant phenotype. Our study identified *pgp-9* as a key player in IVM tolerance and provided functional evidence of its involvement in modulating IVM efficacy in drug-resistant *C. elegans*. Indeed, the genetic deletion of *pgp-9* in the IVR10 resistant background significantly increased IVM efficacy. This role was further validated by RNAi-mediated knock-down in two other independently derived resistant strains, IVR10- 2014 and IVR10-2022.

Importantly, the functional importance of PGP-9 in mediating IVM tolerance was demonstrated using two distinct phenotypic assays: LDA and MA. Interestingly, the extent of restored IVM sensitivity differed between the tests. In the LDA, deletion of *pgp-9* substantially increased IVM sensitivity but did not fully return to N2B levels while it did in the MA. This aligns with previous observations [34], suggesting that *pgp-9* may play a specific non-redundant role in mediating particular physiological responses to IVM, notably those affecting nematode motility. The discrepancy between these assays underscores the complexity of resistance phenotypes and the importance of using complementary approaches when evaluating gene function in drug resistance.

In contrast, deletions of the other *pgp*s had negligeable effects on IVM tolerance, indicating redundant and limited roles in our resistant background. Intriguingly, although *pgp-6* knockdown was previously shown to modulate IVM sensitivity in IVR10 *C. elegans* [31], its deletion had no significant effect in our study, possibly reflecting a context-dependent role.

The contribution of *pgp-9* in IVM tolerance was further supported using a loss-of-function N2B strain, as previously described [26], and through RNAi-mediated knock-down in two additional IVM resistant strains. The effect of RNAi was more moderate compared to the knock-out, likely due to incomplete gene silencing [35]. Overall, these results are partially consistent with previous findings in N2B strains [26], suggesting that multiple *pgp*s may contribute to basal IVM detoxification; however, in our study, only *pgp-9* and to a lesser extent *pgp-3*, were involved in IVM tolerance mechanisms. This implies that the IVM-resistant phenotype may rely on a narrower set of functionally dominant transporters, with *pgp-9* at the centre, compared to a susceptible background.

To assess the evolutionary conservation of PGP-9 function in a parasitic species, we then heterologously expressed *Hco-pgp-9.1* in IVR10(*pgp-9*) worms. This restored the IVM-resistant phenotype, unequivocally demonstrating that *H. contortus* PGP-9.1 can substitute for *C. elegans* PGP-9 thus conferring IVM tolerance. A previous study conducted in mammalian cells demonstrated the ability of heterologous *Hco*-PGP-9.1 to transport Rhodamine 123, a well-known PGP substrate, and its capacity to interact with MLs [18]. Similarly, a yeast-based growth assay indicated that *Pun*-PGP-9 confers protection against ketoconazole toxicity [14], a fungicide also known to interact with PGPs. However, these studies did not inform on the specific contribution PGP-9 to xenobiotic clearance in a drug-resistant strain. Our study builds upon this foundation by providing the first *in vivo* functional demonstration of a conserved detoxification role for a parasitic nematode PGP in a ML-resistant genetic background.

Evidences suggest IVM resistance in nematodes is partly driven by detoxification systems including PGPs, but still IVM dynamics *in vivo* remain poorly understood due to a lack of tools to track the drug at tissue level. To that effect, we developed F-IVM, a fluorescent IVM analog, that closely mimics the parental compound, both structurally and functionally. In N2B animals, F-IVM strongly accumulated in the pharynx and intestine, major sites for xenobiotic uptake and metabolism [36]. In contrast, resistant IVR10 worms showed minimal fluorescent drug accumulation, suggesting enhanced F-IVM efflux or metabolism. This is further supported by a transcriptomic study showing the upregulation of several detoxification genes in IVR10 [25,27], thereby suggesting activation of an IVM detoxification pathway. Strikingly, *pgp-9* deletion restored F-IVM accumulation, particularly at the pharyngeal-intestinal valve and along the intestinal lumen, pointing to disrupted efflux and possible retention at the apical membrane of intestinal epithelial cells in the absence of PGP-9. Quantitative analysis confirmed increased F-IVM uptake in both IVR10(Δ*pgp-9*) and N2B compared to IVR10. However, accumulation patterns in N2B and IVR10(Δ*pgp-9*) did not fully overlap, indicating additional detoxification components are likely involved in IVM clearance [9]. These findings reinforce the idea that PGP-9 acts as a gatekeeper, limiting IVM bioavailability at pharmacologically active tissues.

We then investigated *pgp-9* localization to gain insight on its function in drug clearance. *Pgp-9* localization was consistent with previous studies in *C. elegans* (S4 Table), showing expression in the pharynx and intestine, major sites of drug uptake and detoxification. In *H. contortus*, distinct tissue distribution of one PGP-9 paralog (*e.g.*, uterine expression) suggests paralog-specific physiological roles [18]. Our results are also consistent with another study showing that tissue-specific overexpression of *Pun*-PGP-9 in *C. elegans* confers resistance to MLs in a context-dependent manner [21]. Intestinal expression protected primarily against ingested IVM, while epidermal expression affected drug penetration independently of ingestion. Our study also showed significant *pgp-9* expression in *C. elegans* head neurons, which aligns with scRNA-seq data (S4 Table) and supports the notion of a conserved neuronal detoxification function across nematodes, notably in *P. univalens* [14]. Interestingly, it was previously shown that *C. elegans* and *H. contortus* with dye-filling defects of amphids exhibited drug resistance, and that in a dye-defective *osm-3* mutant, *pgp* overexpression regulated by NHR-8 conferred protecting against Tunicamycin. Therefore, it is likely PGP-9 exerts a protective role against toxic xenobiotics, particularly IVM, in head neurons which are widely enriched in GluCls, the primary IVM target[37]. These findings support a model in which PGP-9 reduces drug entry at physiological barriers, with localization likely influencing its substrate specificity and impact on resistance. Overall, these data support that PGP-9 is located on strategic physiological barriers, and therefore influences IVM accumulation and susceptibility.

## Conclusion

Altogether, our study provides the first *in vivo* functional validation of *pgp-9* causal role in IVM resistance across *C. elegans* and *H. contortus*. We established that *pgp-9* is key to maintain IVM resistance by limiting IVM accumulation at key tissue sites, and thus acting as a detoxifier at strategic physiological barriers of the nematode. Our results show the central role that PGP-9 plays in these nematodes to regulate IVM concentration in tissues where its effects on GluCl receptors induce toxicity to the nematodes. These findings identify PGP-9 as a potential therapeutic target in efforts to counteract IVM resistance, marking a significant step forward in our understanding in resistance mechanisms. Nonetheless, the multifactorial nature of resistance, which likely implicates other PGPs, CYP450s [38], and regulators like NHR-8 [31,32] and CKY-1 [39,40] warrants further investigation. Future work should investigate how these networks interact, whether they operate concomitantly or independently, and determine at which stages of tolerance and resistance development they become critical. Additionally, exploring combinatorial inhibition of these effectors to target parasitic nematodes could offer promising therapeutic strategies.

## Methods

### Materials

All chemicals were obtained from Sigma-Aldrich, unless otherwise stated. IVM and its fluorescent analog, F-IVM, were dissolved in DMSO, and the maximal concentration of DMSO was 0.5% in all assays (except for F-IVM, which was of 1% in the liquid LDA).

### *Caenorhabditis elegans* strains and cultivation conditions

Wild-type *C. elegans* strain N2 Bristol (N2B) and the OP50 *Escherichia coli* strains were provided by the Caenorhabditis Genetics Center (CGC, University of Minnesota, Minnesota, Minneapolis, MN, USA). Mutant strain for *pgp-9* (tm0830), referred as N2B(Δ*pgp-9*), was obtained from the National BioResource Project (Tokyo, Japan). The IVM Resistant strain IVR10 was kindly provided by C. E. James [25].

IVR10-2014 and IVR10-2022 are two IVM-resistant strains that were also selected by step- wise exposure of N2B to increasing concentrations of IVM[31]. IVR10 strains deleted individually for a *pgp* gene, *i.e.*, IVR10; *pgp-1* (knu1129 [Δ7392bp]), IVR10; *pgp-3* (knu1141 [Δ5404bp]), IVR10; *pgp-6* (knu1125 [Δ5208bp]), IVR10; *pgp-9* (knu1140 [Δ9179bp]), IVR10; *pgp-11* (knu1142 [Δ7825bp]) and IVR10; *pgp-13* ((knu1145 [Δ5143bp]) were designed by InVivo Biosystems (Eugene, Oregon, USA). Full length deletions were done in the IVR10 genome using CRISPR-sdm transgenesis method, and a three-frame stop sequence was inserted. In this study, these knock-out strains are referred as IVR10(Δ*pgp-1*), IVR10(Δ*pgp-3*), IVR10(Δ*pgp-6*), IVR10(Δ*pgp-9*), IVR10(Δ*pgp-11*) and IVR10(Δ*pgp-13*). The genotype of each strain was confirmed by PCR and sequencing.

Transgenic strains were generated through microinjection via standard protocols as described in the “Generation of transgenic *C. elegans*” section.

All strains were cultured and handled according to the procedures previously described [27,31]. Nematodes were cultured at 21 °C on Nematode Growth Medium (NGM) agar plates (1.7% bacto agar, 0.2% bacto peptone, 50 mM NaCl, 5 mg/L Cholesterol, 1 mM CaCl_2_, 1 mM MgSO_4_, and 25 mM KPO_4_ Buffer) seeded with *E. coli* strain OP50 as a food source. All *C. elegans* strains were cultured on classic NGM agar plates, except for IVR10, which was cultured on NGM plates containing 10 ng/ml of IVM.

Nematodes were synchronized through egg preparation with sodium hypochlorite. An asynchronous population was collected from NGM plates with M9 buffer (3 g KH2PO4, 6 g Na2HPO4, 5 g NaCl, 0.25 g MgSO4 7H2O in 1 l of water). All larval stages except eggs were lysed with bleaching mix (5 M NaOH, 1% sodium hypochlorite) followed by M9 washes. *C. elegans* eggs were then hatched overnight with mild agitation at 21 °C in M9 to obtain a synchronized population of first-stage larvae (L1).

### Larval development assays (LDAs)

This assay measures the potency of IVM to inhibit the development of *C. elegans* from L1 to young adult. The LDA was essentially conducted as described previously [27,31].

### Solid LDA

Briefly, 30 synchronized L1 larvae were added per well of a 12-well plate. These were seeded on NGM containing increasing concentrations of IVM seeded with OP50 bacteria. Each concentration was set-up in triplicates. Plates were incubated at 21 °C until L1 of the negative control had developed into young adult worms. Development was calculated as a percentage of young adults in the presence of IVM normalized to the untreated control. All experiments were reproduced in at least three biological replicates. Curve fitting for the LDA (sigmoidal dose-response curve with variable slope) was performed with the GraphPad Prism 8.4.2 software. IC_50_ values, the concentrations at which 50% of the animals fail to reach the young adult stage, and the Resistant Factor (RF), the fold resistance relative to N2B were determined.

### Liquid LDA

This assay was performed in liquid media when assessing the ability of an IVM fluorescent analog, F-IVM, to inhibit larval development. In that case, 25 synchronized L1s were seeded in 200 µl of complete liquid S-Basal media seeded with OP50 (5mg/ml). Complete medium was prepared as follows: 50 ml of S-Basal (5.85 g NaCl, 1 g K_2_HPO_4_, 6 g KH_2_PO_4_ in 1 l of water), 500 µl of Potassium Citrate 1 M pH6 (20 g C H O, H O, 293.5 g K C H O, H O in 1 l of water), 500 µl of Trace Metal Solution (1.86 g C H N Na O, 0.69 g FeSO_4_, 7H_2_O, 0.2 g MnCl_2_, 4H_2_O, 0.29 g ZnSO_4_, 7H_2_O, 0.025g CuSO_4_, 5H_2_O), 150 µl CaCl_2_ 1 mM, 150 µl MgSO_4_ 1 mM, 50 µl Cholesterol 5 mg/ml. Drug treatment was administered by adding 1-2 µl of F-IVM at increasing concentrations. Plates were incubated at 21 °C under gentle agitation until L1 of the DMSO control had developed into young adult worms. Development, dose-response curves, IC_50_s and RF were determined as described above.

### RNA interference on *pgp-9* in IVR10 strains

RNA interference (RNAi) was conducted by feeding HT115 bacteria transformed with L4440 vector that produces double-stranded RNA against a targeted gene to the strains, as previously described[31]. HT115 bacteria clones expressing *pgp-9* RNAi or the empty vector as control from the Ahringer RNAi library were grown for 8h at 37 °C in LB medium containing ampicillin (50 μg/ml). They were then seeded on NGM plates supplemented with carbenicilin (25 μg/ml) and IPTG (1 mM) to induce RNAi expression. Finally, solid LDA was conducted as described above. The extent of knockdown of *pgp-9* mRNA was determined by RT-qPCR.

### LDA for transgenic strains

This assay was adapted from another study [41] and from the solid LDA previously described to characterize the rescue function of *Hco-pgp-9.1* in the IVR10 *pgp-9* knock-out strain. Synchronized L1 larvae were obtained from bleaching a population heterogeneously carrying the *Hco-pgp-9.1* extrachromosomal array. 60 L1s were then added per well of a 6-well plate seeded with NGM and OP50, and containing increasing concentrations of IVM, from 0 (*i.e.*, DMSO) to 10 nM. Each concentration was set-up in sextuplicate. Plates were incubated at 21 °C until L1 of the DMSO control had developed into young adult worms. The number of worms negative and positive for the extrachromosomal array, respectively IVR10(Δ*pgp-9*) and the *Hco*-PGP-9 rescue strain, were then counted in each well. The percentage of animals reaching the adult stage for each strain was expressed as a fraction normalized to the number of adult worms of each population present in the DMSO well. All experiments were reproduced in biological triplicate, and three independent transgenic strains of the *Hco*-PGP-9.1 rescue were studied.

### Motility assay (MA)

This assay measures the potency of IVM to inhibit the motility of *C. elegans* young adults. The MA was essentially conducted as described previously, using the WMicroTracker™ One device (PhylumTech, Santa Fe, Argentina) [34]. Synchronized young adults (40 per well) were seeded into 200 µl of M9 in a 96-flat well plate. Before each measurement, plates were incubated 15 min at 21 °C to allow the worms to settle. Basal activity (BA) was then measured for 30 min to normalize movement activity in each well. Drug treatment was administered by adding 1µl of IVM at increasing concentrations. Following drug treatment, score activity (SA) was recorded for a 120 min period. Negative controls (NG), *i.e.*, wells without worms were conducted. Motility percentages were calculated for each treated well as fold induction relative to DMSO treated worms which was set to 100, *i.e.*, SA-NC_120min_/BA- NC_30min_. All experiments were reproduced in technical triplicates and at least three biological replicates. Curve fitting for the MA (sigmoidal dose-response curve with variable slope), IC_50_ values, and RFs were determined as explained above.

### RT-qPCR

A population of 1500 synchronized L1 larvae were added to NGM plates seeded with HT115 or HT115-*pgp-9* bacteria in order to assess RNAi efficiency. Non-gravid young adults were then collected using M9 buffer and flashed frozen in liquid nitrogen in RLT buffer (Qiagen) supplemented with DTT 2 M. Lysed worms were stored -80 C. Frozen samples were thawed and homogenized three times for 10 sec at 6 m.s^−1^ in a FastPrep-24 instrument (MP-Biomedicals, NY, USA). Total RNA was extracted using RNeasy Plus Kit (Qiagen, S.A., Courtaboeuf, France) according to the manufacturer’s instructions. Total RNA was quantified using a NanoDrop ND-1000 spectrophotometer (NanoDrop Technologies Inc., Wilmington, DE, USA). cDNA was synthesized from 1 μg of total RNA using Maxima H Minus First Strand cDNA Synthesis Kit (Thermofisher).

Real-time quantitative polymerase chain reaction (RT-qPCR) was performed using SYBR™ Green PCR Master Mix (Applied BiosystemsLife Technologies, Courtaboeuf, France) and a CFX96 Touch Real-Time PCR Detection System (Bio-Rad). Gene specific primers that were used are listed in S1 Table. Results are expressed according to the relative quantification method with *tba-1* as the reference gene.

### Identity between *Cel*-PGP-9 and *Hco*-PGP-9.1

Predicted amino acid sequences of *Cel*-PGP-9 (Accession Number: CE15714) and *Hco*-PGP-9.1 (Accession Number: HCON_00130050-00001) were obtained from WormBase and WormParasite, respectively. Alignment was first performed with full-length sequences, with the Clustal Omega Multiple Sequence Alignment (MSA) tool (https://www.ebi.ac.uk/jdispatcher/msa/clustalo?stype=protein). Percent Identity Matrices were extracted following the alignment. Nucleotide Binding Domains (NBDs) were then identified with the help of the Scan Prosite tool (https://prosite.expasy.org/scanprosite/) (and then manually curated from the sequences such as: *Cel*-PGP9 NBD1(383-619); NBD2(1030-1294) and *Hco-*PGP-9.1 NBD1(373-609); NBD2(1026-1270). Sequences were then subjected again to Clustal Omega and new identity matrices were obtained.

### Generation of transgenic *C. elegans*

#### Cloning

Unless stated otherwise, all PCRs were performed with the proofreading Phusion High-Fidelity DNA Polymerase (New England Bioloabs, Ipswich, MA, USA) and all cloning were carried out with the Pro Ligation-Free Cloning Kit (abm, Richmond, BC, Canada), following the manufacturer’s information. Primers used to design the plasmids are listed in S2 Table. A 2.6 kilo base pairs region was amplified from N2B genomic DNA to isolate *pCel-pgp-9* and was subcloned into the pMini T2.0 vector (New England BioLabs). This promoter region was then cloned into a PstI/SmaI digested plasmid containing fused *SL2*::*mCherry*::*Cel-unc-54*.

*Hco-pgp-9.1* originated from the subcloning vector pGEM-T Easy from Pr. Roger Prichard team (McGill University, Parasitology Institute)[18]. It was cloned into the plasmid containing *pCel-pgp-9*::*SL2*::*mCherry*::*Cel-unc-54*, between *pCel-pgp9* and *SL2* (restriction sites SmaI/KpnI) (GeneCust). Plasmid constructs were systematically verified by sequencing (Eurofins).

#### Microinjections

Plasmid constructs, i.e, *pCel-pgp-9::SL2::mCh::Cel-unc-54* and *pCel-pgp- 9::Hco-pgp-9.1::SL2::mCh::Cel-unc-54*, along with co-injection plasmid: pSC391, expressing YFP in neurons, (a gift from Dr. David Miller, Vanderbilt University) or pPD118.3, expressing GFP in the pharynx, (a gift from SEGiCel, SFR Santé Lyon Est CNRS UAR 3453, Lyon, France), were microinjected into either N2B(Δ*pgp-9*) or IVR10(Δ*pgp-9*) gonads (Institute of Parasitology, McGill University, Canada or SEGiCel, SFR Santé Lyon Est CNRS UAR 3453, Lyon, France).

Worms were prepared as described elsewhere [42]. Young adult hermaphrodites N2B(Δ*pgp-9*) were transformed by microinjection of *pCel-pgp-9::SL2::mCh::Cel-unc-54* at 75 ng/μL and pSC391 at 15 ng/µl (3:1 ratio) into the gonads using an Eppendorf Femtojet 4 connected to an Eppendorf InjectMan NI 2 (Institute of Parasitology, McGill University, Canada) and homemade needles (MODEL P-1000, Sutter Instrument).

IVR10(Δ*pgp-9*) were transformed with either *pCel-pgp-9::SL2::mCh::Cel-unc-54* or *pCel-pgp-9::Hco-pgp-9.1::SL2::mCh::Cel-unc-54* at 50 ng/µl and pPD118.3 at 5 ng/µl. DNA concentrations were adjusted up to 120 ng/µl using a 1kb Plus DNA ladder (Invitrogen). Microinjections were carried out at SEGiCel (Claude Bernard Lyon 1 University, France) using a General Valve Corporation Picospritzer II Microinjector/Transjector and homemade needles (Narishige PC-10 Dual Stage Glass Micropipette Puller).

Successful transformations were identified using an SMZ800N fluorescent stereomicroscope (Nikon). At least three individual strain lines carrying extrachromosomal arrays were obtained for each construct.

Transgenic strains were maintained by picking regularly GFP or mCh-fluorescent individuals to a new plate, as described in the “*Caenorhabditis elegans* strains and cultivation conditions” section. Development assays were conducted on the IVR10(Δ*pgp-9*) carrying *Hco-pgp-9.1* in an extrachromosomal array. Three independent strains were studied and are referred to as: IVR10(Δ*pgp-9*)-R#1, IVR10(Δ*pgp-9*)-R#2, and IVR10(Δ*pgp-9*)-R#3. Transcription of *Hco-pgp-9.1* was confirmed by single worm RT-qPCR. RNA extraction was conducted as described elsewhere [43] and qPCR was performed as explained above.

### Fluorescent IVM (F-IVM) probe

#### Synthesis of F-IVM

IVM (6 mg, equivalent to 150 µL of IVM diluted to 40 mg/mL in methanol (MeOH)) was mixed with 100 µL of 1-N-methyl-imidazole (MI), followed by sequential additions of 150 µL of anhydrous trifluoroacetic acid (TFA) and 50 µL of triethylamine (TEA), with vortexing after each addition. The reaction mixture was incubated at 70°C for 1 h. The resulting F-IVM was purified using solid phase extraction (SPE) on LC18-100 mg cartridges with an automated SPE robot (Rapidtrace). Cartridges were conditioned with 1 mL MeOH (5 min, 0.2 mL/sec) followed by 1.2 mL H2O (6 min, 0.2 mL/sec). Samples were loaded twice onto cartridges at 0.1 mL/sec for 4.2 min and 4 min, respectively, and left to stand for 1 min. The cartridges were washed sequentially with 2 mL H2O (0.1 mL/sec) and 1 mL MeOH/H2O (25/75) at 0.2 mL/sec, then dried with an air stream (10 min, 0.5 mL/sec). F-IVM was eluted with acetonitrile (AcN, 0.5 mL/sec) and dried under a nitrogen stream (N2) at 60°C. A portion of the eluates was analyzed using an HPLC-Fluorescence system (Ultimate 3000 Thermo, ThermoFisher, Waltham, MA, USA). Samples (25 µL) were injected onto a Supelcosil LC-18 column (150 x 4.6 mm; 3 µm Supelco) under isocratic elution with a mobile phase of H2O (0.4% of acetic acid)/MeOH/AcN (5/30/65, v/v/v) at a flow rate of 1.6 mL/min. The fluorescence detector was set to an excitation wavelength of 355 nm and an emission wavelength of 465 nm. Column temperature was maintained at 40°C, and the autosampler was kept at 20°C.

### Characterization of F-IVM

The structure of F-IVM was confirmed by comparison to native IVM using mass spectrometry (MS). MS analyses were performed in MS scan and daughter scan modes on a triple quadrupole instrument (Xevo TQ, Waters, Milford, MA, USA) with electrospray ionization in positive mode (ESI+). Key parameters included a capillary voltage of 3.5 kV, source temperature of 150 °C, desolvation temperature of 600 °C, and cone voltage of 25 V. Fragmentation was induced using argon as collision gas (collision energy: 20 eV).

### F-IVM stability study

Short-term stability of the F-IVM probe was assessed at room temperature (24 and 48 h) and 4 °C (24 h, 48 h, and 12 days). Long-term stability was evaluated through repeated freeze–thaw cycles and continuous storage at –80 °C for up to one year. Quantification was performed by HPLC-fluorescence as previously described, using a fluorescent ivermectin derivative (IVM-TA) as an external standard. Stability was determined by comparing F-IVM/IVM-TA area ratios to initial values (T).

### Confocal microscopy

#### Transgenic strain imaging

An asynchronous population of transgenic worms was washed once with M9 buffer to remove OP50 bacteria. Worms were paralyzed with 2.5 mM Sodium Azide and mounted on glass. mCherry, expressed under the native promoter of *Cel-pgp-9* in N2B and IVR10 *pgp-9* knock-outs, was visualized with Zeiss LSM 710 AxioObserver Plan-Apochromat 20x/0.8 M27 or 40x/1.3 Oil Iris (Infinity-INSERM UMR1291, France). mCh was excited at 561 nm and recorded between 581 and 664 nm, and brightfield images were recorded to aid anatomical interpretation.

### F-IVM quantification

Accumulation of the fluorescent F-IVM was investigated in adult N2B, IVR10, and IVR10(Δ*pgp-9*) *C. elegans* that were grown in liquid media. Synchronized L1s were incubated in complete liquid media S-Basal seeded with OP50 at 21 °C under gentle agitation. Complete medium was prepared as explained above. After a 72 h growth period, animals were collected and then incubated with F-IVM at 10 µg/ml in complete liquid media seeded with OP50 at 2 1°C under gentle agitation. After 72 h, they were washed three times with M9 buffer to remove F-IVM and incubated for 4 hours in M9 buffer. Before imaging, samples were once again washed, and worms were paralyzed using Levamisole 2.5 mM and mounted on glass. F-IVM fluorescence was visualized and images were acquired with a Zeiss LSM 710 AxioObserver Plan-Apochromat 20x/0.8 M27 (Multi-Scale Imaging Facility, McGill University, Canada). F-IVM fluorescence was recorder as follows: diode laser 405 2%, emission wavelength 482 nm and detection wavelengths 414-550 nm with a detector gain of 620. Auto-fluorescence was recorded on worms that were not incubated with F-IVM. Images were acquired using ZEN 2.3 software. For each sample, the Mean Fluorescence Intensity, meaning the Integrated Intensity (A.U.) divided by the worm area (pixel²), was evaluated. It was then normalized with auto-fluorescence, *i.e.*, the Mean Fluorescence Intensity of strains unlabeled with F-IVM. Experiments were replicated three times, and approximately 20-30 worms were analyzed per condition.

### Statistical analysis

All experiments were conducted independently at least in triplicate, except for the F-IVM MA which was in duplicate. Results are expressed as mean ± standard deviation (S.D.) or ± standard error of the mean (S.E.M.). Statistical analyses were performed using the GraphPad Prism 8.4.2 software, and results were considered statistically significant when p < 0.05.

To evaluate the impact of each *pgp* deletion in the IVR10 background compared to the control background strain IVR10, a one-way analysis of variance (ANOVA) followed by Dunnett’s multiple comparisons test was performed on the knock-out or N2B strain compared to IVR10. Only key statistical results are presented in the study to maintain clarity and avoid excessive details. To assess the effect of *pgp-9* invalidation in the N2B strain and of *pgp-9* silencing in individual IVM-resistant strains, unpaired parametric t-tests were performed.

The expressions of *Hco-pgp-9.1* in the three transgenic strains were individually compared to that of *Cel-pgp-9* in IVR10 using unpaired parametric t-tests. F-IVM accumulation in N2B, IVR10 and IVR10(Δ*pgp-9*) was statistically assessed with a one-way ANOVA followed by a Tukey’s post-hoc test. Finally, LDAs with transgenic strains were statistically analyzed using a two-way ANOVA to assess the effects of the IVM concentrations and *Hco-pgp-9.1* expression in IVR10(Δ*pgp-9*) on larval development. Pairwise comparisons between IVR10(Δ*pgp-9*) and IVR10(Δ*pgp-9*) expressing the transgene at each concentration were performed using Sidak’s multiple comparisons test. The LDAs of each of the three transgenic strains were analyzed individually.

## Acknowledgments

We thank Rémy Betous for his support and advice in Molecular Biology, and Lucas Barat for the numerous scientific discussions and his punctual assistance. We would like to thank the McGill Institute of Parasitology Institute and the Multi-Scale Imaging Facility where one of the rescue models and part of the confocal images were respectively performed. We also thank Hua Che for the technical support at McGill and Cody Hart, who facilitated the acquisition of microinjection skills. We thank Agreenium, DESSE, and INRAE for the scholarship they granted for a doctoral mobility at McGill University. The rest of the transgenic strains were generated at SEGiCel (SFR Santé Lyon Est CNRS UAR 3453, Lyon, France) with the support of CNRS and IBiSA. We thank Thomas Boulin, Margaux Gibert, and Marielle Limoges for their support. Some of the confocal imaging experiments were also performed at the Infinity-INSERM UMR1291 core facility connected to the Toulouse Réseau Imagerie network, member of the France-BioImaging national infrastructure supported by the French National Research Agency (ANR-10-INBS-04). For that, we thank Simon Lachambre for his assistance with the confocal imaging systems. We also thank Cécile Pouzet for the technical assistance at the TRI-FRAIB Imaging Facility (Université de Toulouse-CNRS). This work was supported by grant N° ANR-21-CES35-004 “Fluo-RES” from the Agence Nationale de la Recherche.

## Authors contribution

Conceptualization: C.B., M.A., A.L.

Data curation: C.B., M.A., M.L.

Formal analysis: C.B., M.A., M.L.

Funding acquisition: M.A., A.L.

Investigation: C.B., M.A., E.G, J.F.S., F.R.P.

Methodology: C.B., M.A., E.C.

Supervision: M.A., A.L., M.L.

Writing – original daft preparation: C.B.

Writing – review & editing: C.B., M.A., A.L., R.P., M.L., E.C.

## Data availability

Source data are provided at: https://entrepot.recherche.data.gouv.fr/dataset.xhtml?persistentId=doi:10.57745/HP17LZ

## Supplementary figures

**S1 Figure.**
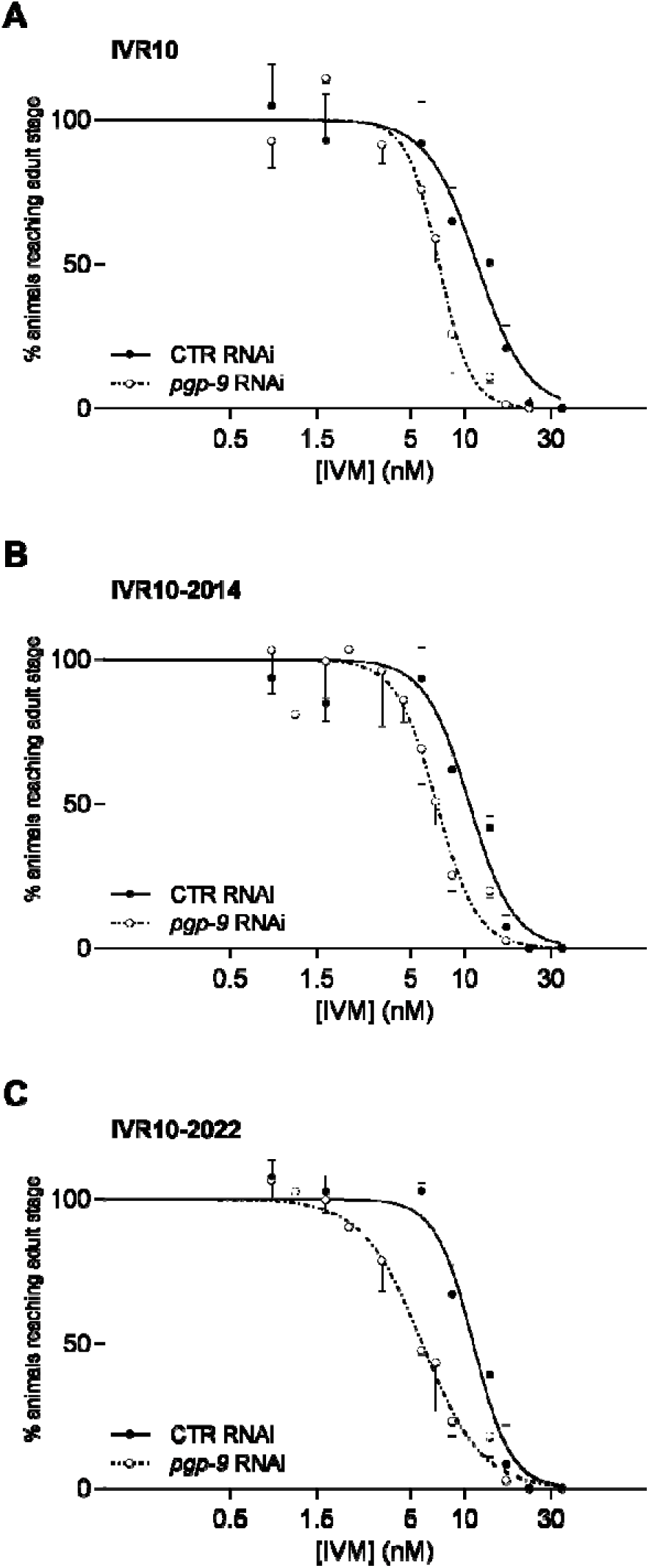
*pgp-9* silencing in IVM-resistant strains. Effect of *pgp-9* silencing on susceptibilities of IVM-resistant strains IVR10 (**A**), IVR10-2014 (**B**) and IVR10-2022 (**C**) to ivermectin (IVM) in a larval development assay (LDA). Worms were fed on bacteria carrying a plasmid producing double stranded RNA against *pgp-9* plasmid (*pgp-9* RNAi) or an empty plasmid as control (CTR RNAi). Values of dose-response curves represent the percentage of young adults maintaining motility within the presence of increasing doses of IVM. Data are mean ± S.D. from 3 independent experiments. IC_50_s for each strain are presented in S3 Table. The efficiency of *pgp-9* knock-down was assessed by RT-qPCR and is represented in S1 Figure.

**S2 Figure.**
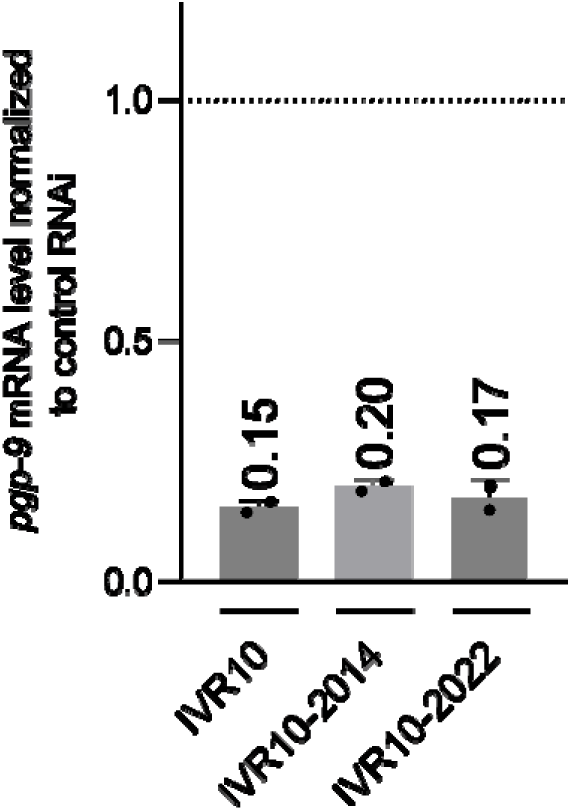
*pgp-9* mRNA expression following silencing. Quantification of *pgp-9* transcripts in IVR10, IVR10-2014, and IVR10-2022 following gene silencing. Real-time RT-qPCR analysis was applied after RNAi treatment with control RNAi or specific *pgp-9* RNAi. *pgp-9* mRNA levels in IVR10 strains are expressed as fold change relative to control RNAi. Data were normalized against *tba-1* as an internal control and are mean ± S.D. from two independent RNA preparations for each strain.

**S3 Figure.**
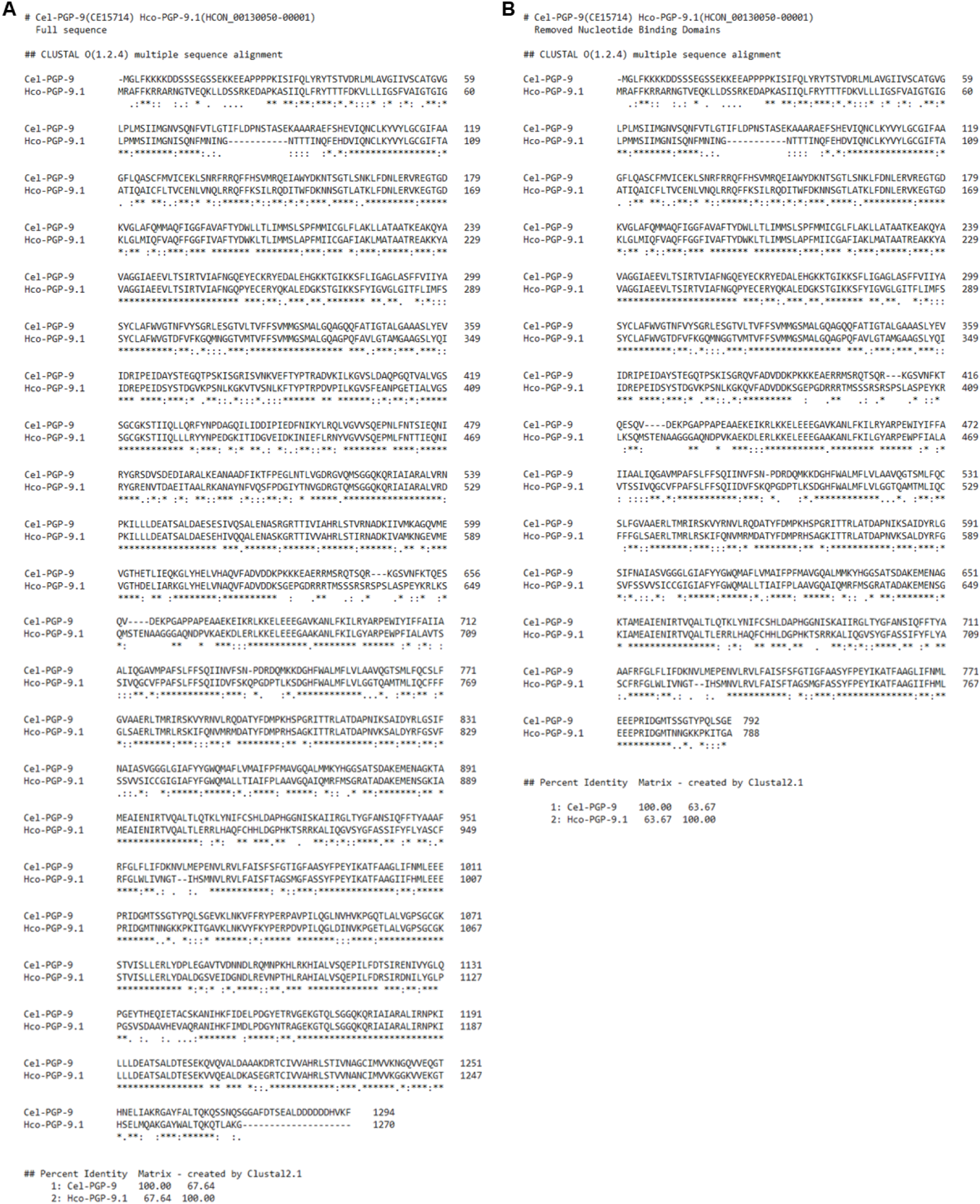
Identity between *Cel*-PGP-9 and *Hco*-PGP-9.1. Sequence alignments and percent identity matrices between *Cel-*PGP-9 (Accession Number: CE15714) and *Hco-*PGP-9.1 (Accession Number: HCON_00130050-00001) were conducted with the Clustal Omega tool. Symbols indicate: (*) conserved amino acid; (:) strong conservation following a substitution; (.) low similarity following a substitution; ( ) no conservation. (**A**) Alignment was first performed with full predicted sequences and the percentage of identity between the two proteins was extracted. (**B**) Nucleotide binding domains (NBDs) were identified with the Scan Prosite tool and then manually curated from the sequences. Sequences were subjected a second time to Clustal Omega in order to highlight homology related to the substrate catalytic part of the protein (*i.e.*, the transmembrane domains (TMDs).

**S4 Figure.**
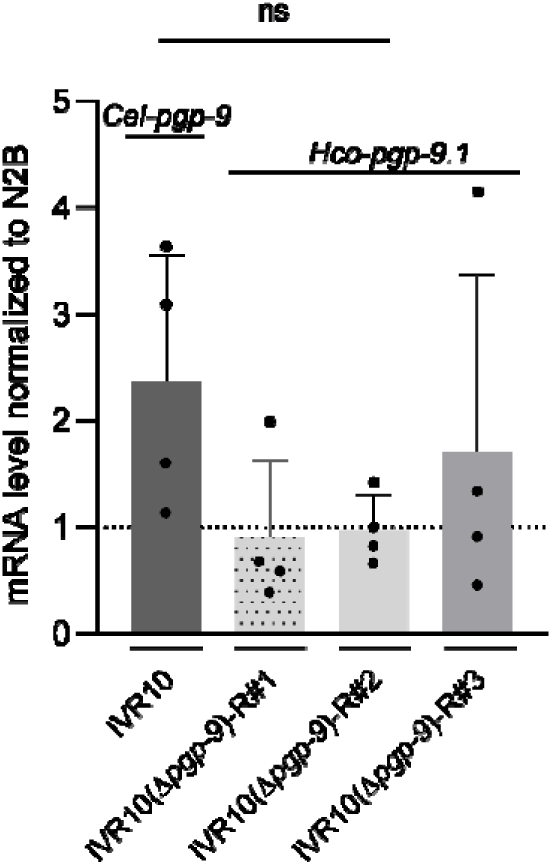
Expression of *Hco-pgp-9.1* transgene. Quantification of *Cel-pgp-9* in IVR10 and *Hco-pgp-9.1* in the transgenic rescue strains, *i.e.*, IVR10(Δ*pgp-9*)-R#1, #2 and #3, by single worm RT-qPCR. Data are expressed as fold change to the expression level of *Cel-pgp-9* in the wild-type strain N2B. *pgp-9* mRNA levels were normalized against the housekeeping gene *tba-1* and are mean ± S.D. from four independent mRNA preparations per strain. One independent mRNA preparation corresponds to an RNA extraction from one single worm. mRNA levels of *Hco-pgp9.1* were compared to those of the IVR10 background strain as a reference (unpaired parametric t-test, ns).

## Supplementary tables

**S1 Table.**
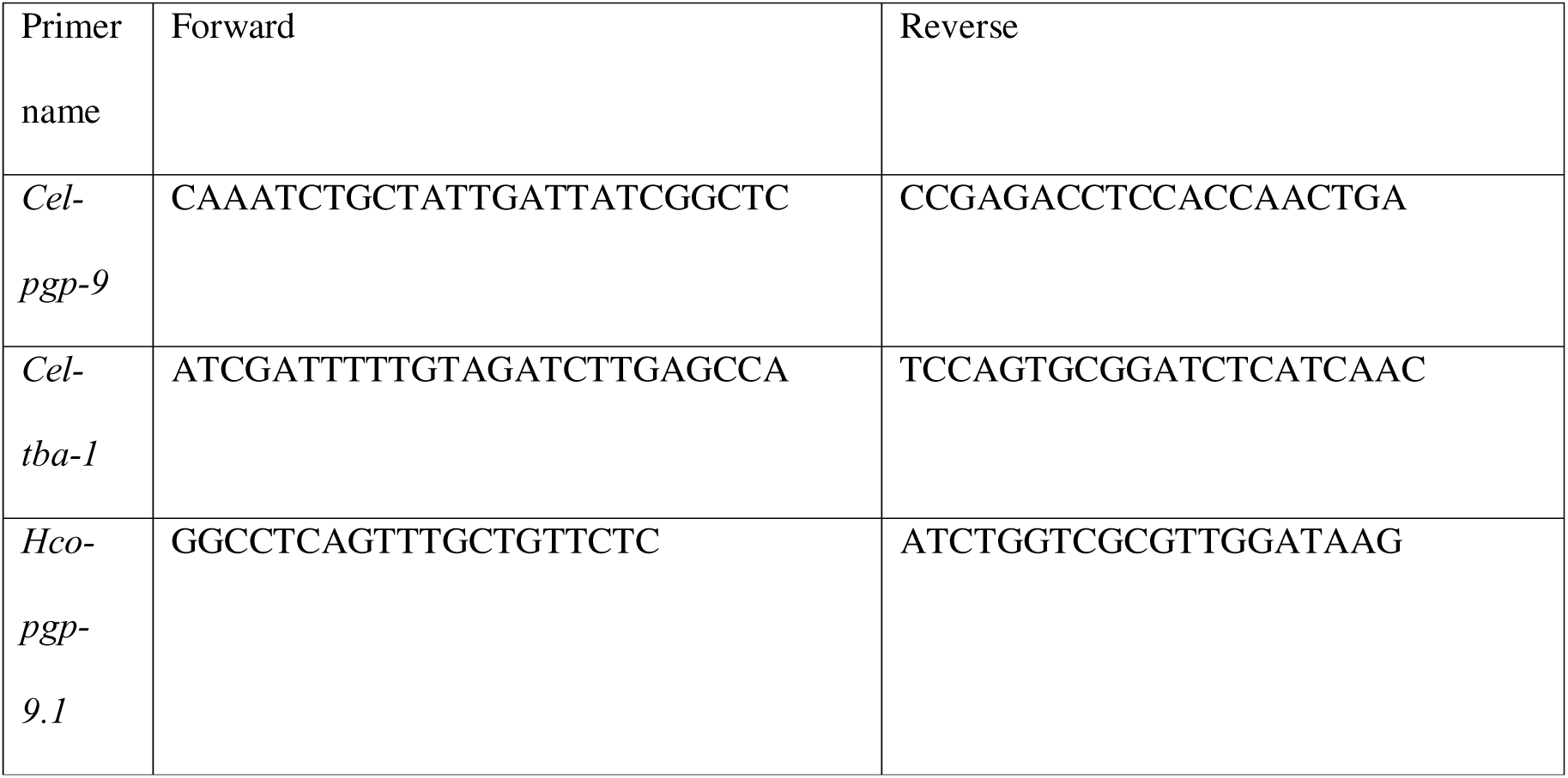
Specific primers for *Caenorhabditis elegans* or *Haemonchus contortus* genes targeted by RT-qPCR.

**S2 Table.**
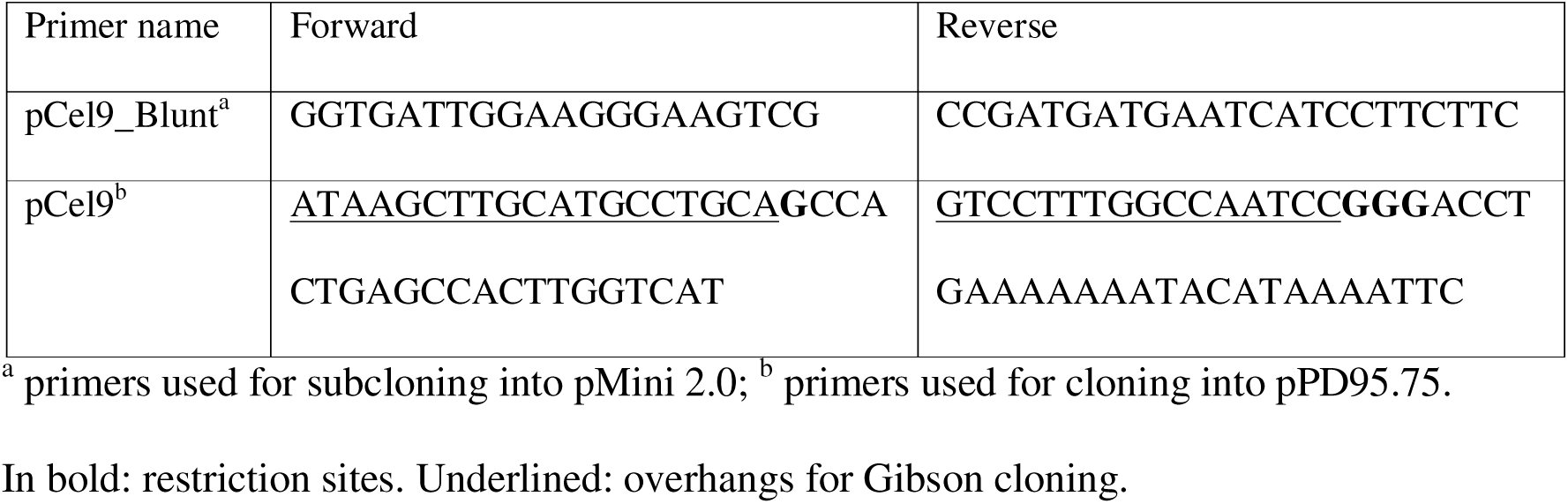
Primers used for cloning in this study.

**S3 Table.**
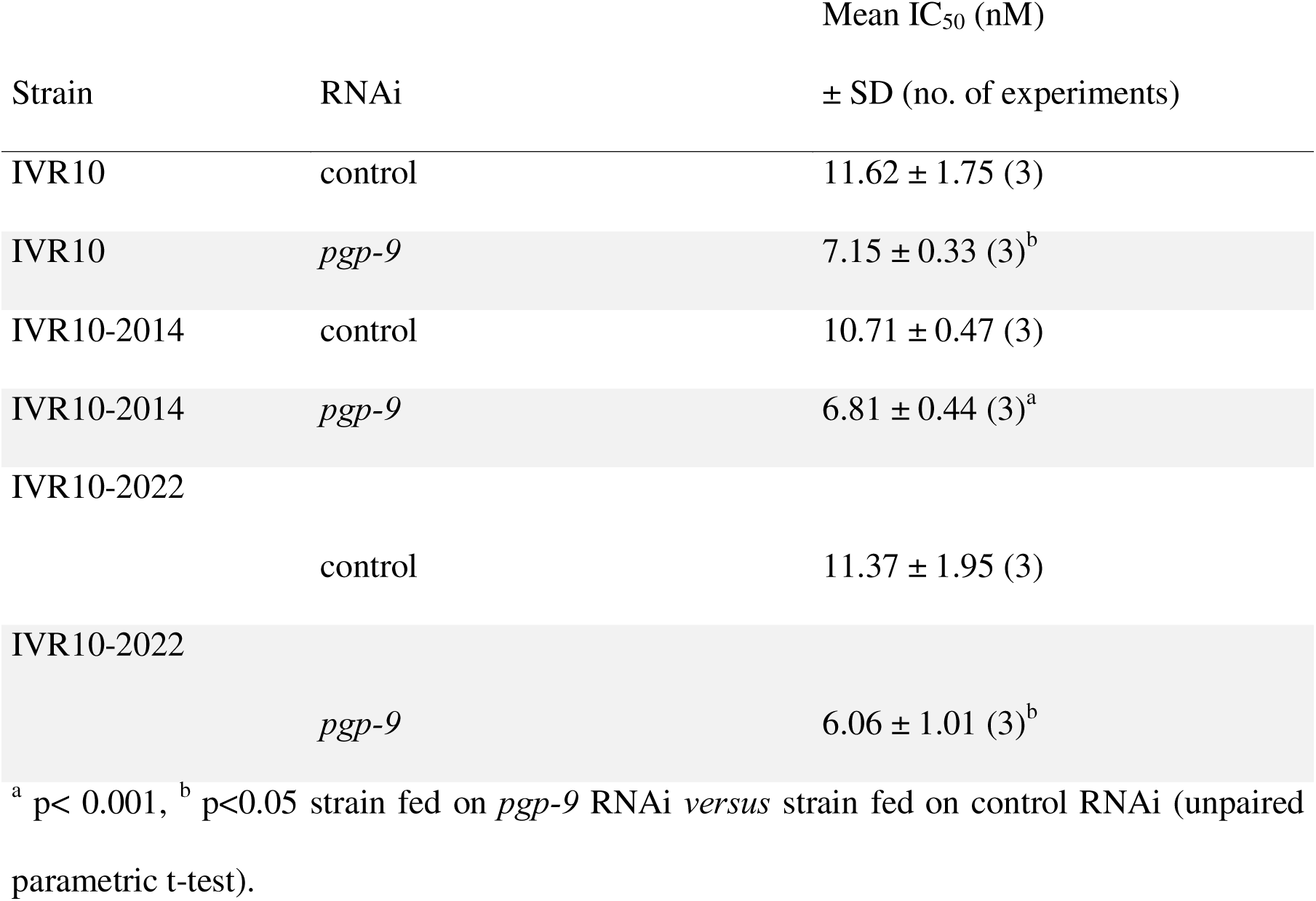
Susceptibilities to ivermectin (IVM) of IVM selected strains IVR10, IVR10-2014, and IVR10-2022 following *pgp-9* silencing on larval development assay (LDA). IC_50_: inhibitory concentration 50%.

**S4 Table.**
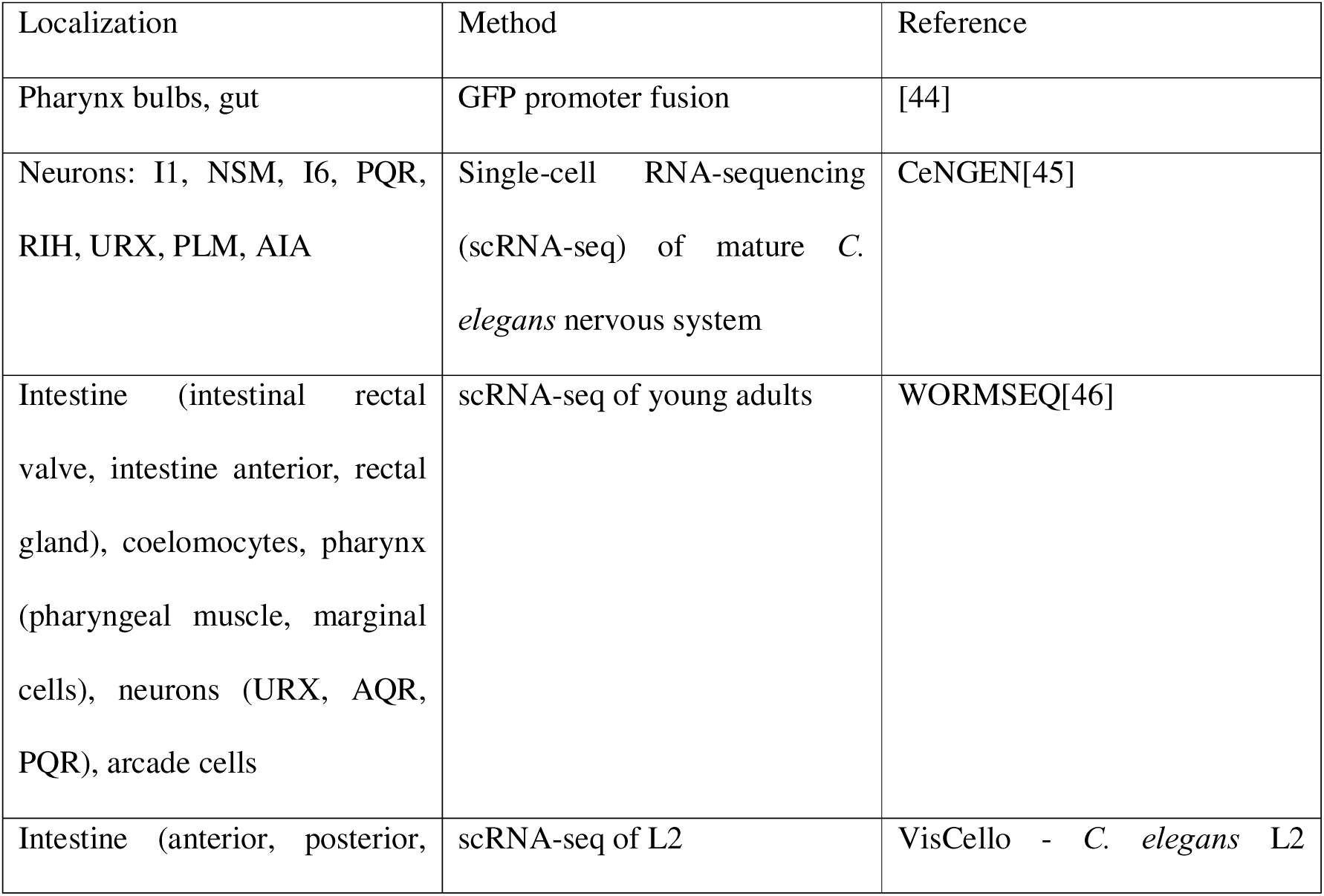
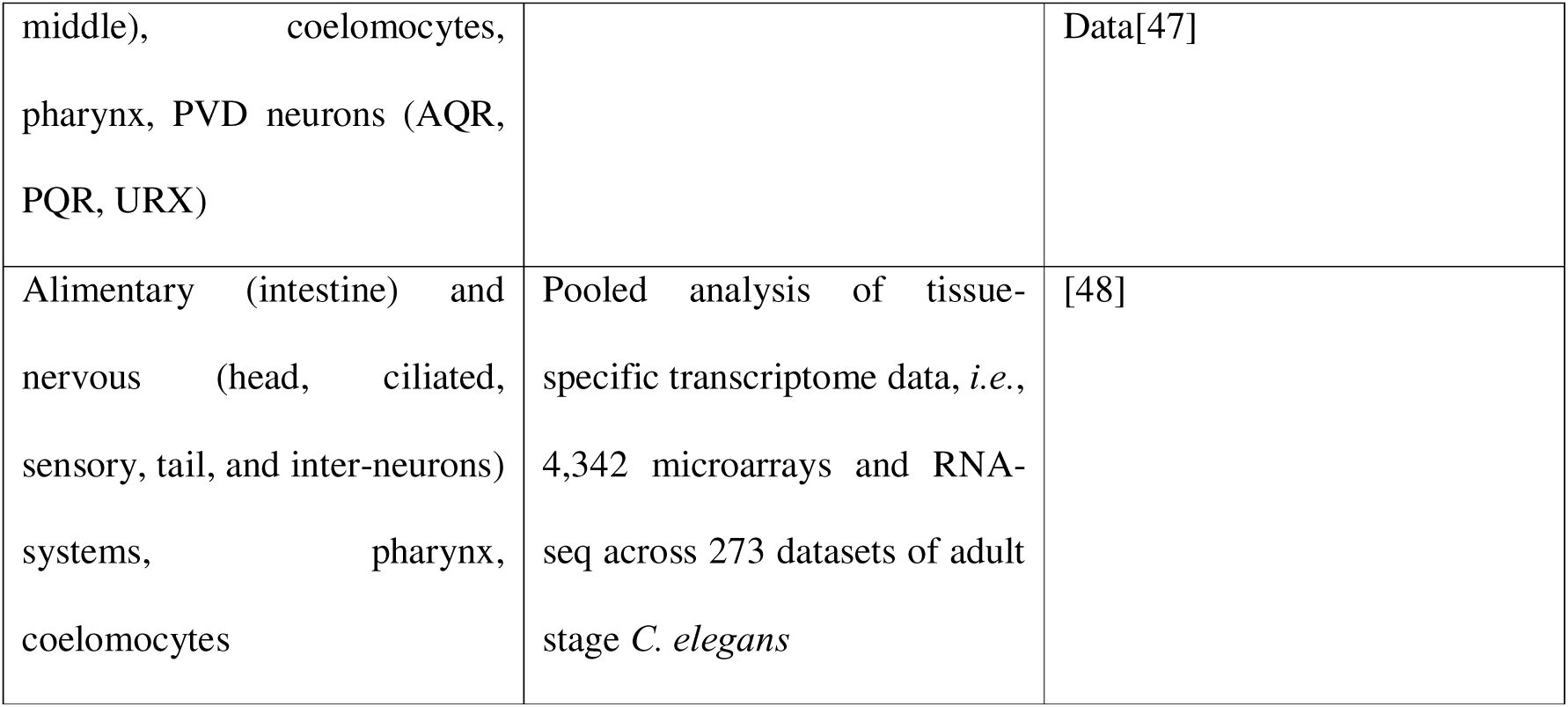
Evidence of *Cel-pgp-9*/PGP-9 localization from the literature.

## References

1. Specht S, Keiser J. Helminth infections: Enabling the World Health Organization Road Map. International Journal for Parasitology. 2023;53: 411–414. doi:10.1016/j.ijpara.2022.10.006

2. Anisuzzaman null, Tsuji N. Schistosomiasis and hookworm infection in humans: Disease burden, pathobiology and anthelmintic vaccines. Parasitol Int. 2020;75: 102051. doi:10.1016/j.parint.2020.102051

3. Zajac AM, Garza J. Biology, Epidemiology, and Control of Gastrointestinal Nematodes of Small Ruminants. Vet Clin North Am Food Anim Pract. 2020;36: 73–87. doi:10.1016/j.cvfa.2019.12.005

4. Charlier J, Rinaldi L, Musella V, Ploeger HW, Chartier C, Vineer HR, et al. Initial assessment of the economic burden of major parasitic helminth infections to the ruminant livestock industry in Europe. Preventive Veterinary Medicine. 2020;182: 105103. doi:10.1016/j.prevetmed.2020.105103

5. Martin RJ, Robertson AP, Choudhary S. Ivermectin: An Anthelmintic, an Insecticide, and Much More. Trends in Parasitology. 2021;37: 48–64. doi:10.1016/j.pt.2020.10.005

6. Kotze AC, Prichard RK. Anthelmintic Resistance in *Haemonchus contortus*. In: Gasser RB, Samson-Himmelstjerna GV, editors. Advances in Parasitology. Academic Press; 2016. pp. 397–428. doi:10.1016/bs.apar.2016.02.012

7. Lustigman S, McCarter JP. Ivermectin Resistance in Onchocerca volvulus: Toward a Genetic Basis. PLOS Neglected Tropical Diseases. 2007;1: e76. doi:10.1371/journal.pntd.0000076

8. Charlier J, Thamsborg SM, Bartley DJ, Skuce PJ, Kenyon F, Geurden T, et al. Mind the gaps in research on the control of gastrointestinal nematodes of farmed ruminants and pigs. Transboundary and Emerging Diseases. 2018;65: 217–234. doi:10.1111/tbed.12707

9. Lespine A, Blancfuney C, Prichard R, Alberich M. P-glycoproteins in anthelmintic safety, efficacy, and resistance: (Trends in Parasitology 40, 896–913; 2024). Trends in Parasitology. 2025;41: 334. doi:10.1016/j.pt.2025.02.003

10. Raza A, Kopp SR, Bagnall NH, Jabbar A, Kotze AC. Effects of in vitro exposure to ivermectin and levamisole on the expression patterns of ABC transporters in Haemonchus contortus larvae. International Journal for Parasitology: Drugs and Drug Resistance. 2016;6: 103–115. doi:10.1016/j.ijpddr.2016.03.001

11. Pacheco PA, Louvandini H, Giglioti R, Wedy BCR, Ribeiro JC, Verissimo CJ, et al. Phytochemical modulation of P-Glycoprotein and its gene expression in an ivermectin-resistant Haemonchus contortus isolate in vitro. Veterinary Parasitology. 2022;305: 109713. doi:10.1016/j.vetpar.2022.109713

12. Tuersong W, Zhou C, Wu S, Qin P, Wang C, Di W, et al. Comparative analysis on transcriptomics of ivermectin resistant and susceptible strains of Haemonchus contortus. Parasites Vectors. 2022;15: 159. doi:10.1186/s13071-022-05274-y

13. Mate L, Ballent M, Cantón C, Lanusse C, Ceballos L, Alvarez L LI, et al. ABC-transporter gene expression in ivermectin-susceptible and resistant Haemonchus contortus isolates. Veterinary Parasitology. 2022;302: 109647. doi:10.1016/j.vetpar.2022.109647

14. Gerhard AP, Krücken J, Heitlinger E, Janssen IJI, Basiaga M, Kornaś S, et al. The P- glycoprotein repertoire of the equine parasitic nematode Parascaris univalens. Sci Rep. 2020;10: 13586. doi:10.1038/s41598-020-70529-6

15. Choi Y-J, Bisset SA, Doyle SR, Hallsworth-Pepin K, Martin J, Grant WN, et al. Genomic introgression mapping of field-derived multiple-anthelmintic resistance in Teladorsagia circumcincta. Andersen EC, editor. PLoS Genet. 2017;13: e1006857. doi:10.1371/journal.pgen.1006857

16. Mealey KL. Therapeutic implications of the MDR-1 gene. J Vet Pharmacol Ther. 2004;27: 257–264. doi:10.1111/j.1365-2885.2004.00607.x

17. David M, Lebrun C, Duguet T, Talmont F, Beech R, Orlowski S, et al. Structural model, functional modulation by ivermectin and tissue localization of Haemonchus contortus P- glycoprotein-13. International Journal for Parasitology: Drugs and Drug Resistance. 2018;8: 145–157. doi:10.1016/j.ijpddr.2018.02.001

18. Godoy P, Che H, Beech RN, Prichard RK. Characterisation of P-glycoprotein-9.1 in Haemonchus contortus. Parasites Vectors. 2016;9: 52. doi:10.1186/s13071-016-1317-8

19. Mani T, Bourguinat C, Keller K, Ashraf S, Blagburn B, Prichard RK. Interaction of macrocyclic lactones with a Dirofilaria immitis P-glycoprotein. International Journal for Parasitology. 2016;46: 631–640. doi:10.1016/j.ijpara.2016.04.004

20. Jesudoss Chelladurai JRJ, Jones DE, Brewer MT. Characterization of a P-glycoprotein drug transporter from Toxocara canis with a novel pharmacological profile. International Journal for Parasitology: Drugs and Drug Resistance. 2021;17: 191–203. doi:10.1016/j.ijpddr.2021.10.002

21. Gerhard AP, Krücken J, Neveu C, Charvet CL, Harmache A, von Samson-Himmelstjerna G. Pharyngeal Pumping and Tissue-Specific Transgenic P-Glycoprotein Expression Influence Macrocyclic Lactone Susceptibility in Caenorhabditis elegans. Pharmaceuticals. 2021;14: 153. doi:10.3390/ph14020153

22. Janssen IJI, Krücken J, Demeler J, von Samson-Himmelstjerna G. Transgenically expressed Parascaris P-glycoprotein-11 can modulate ivermectin susceptibility in Caenorhabditis elegans. International Journal for Parasitology: Drugs and Drug Resistance. 2015;5: 44–47. doi:10.1016/j.ijpddr.2015.03.003

23. Raza A, Kopp SR, Jabbar A, Kotze AC. Effects of third generation P-glycoprotein inhibitors on the sensitivity of drug-resistant and -susceptible isolates of Haemonchus contortus to anthelmintics in vitro. Veterinary Parasitology. 2015;211: 80–88. doi:10.1016/j.vetpar.2015.04.025

24. Jesudoss Chelladurai JRJ, Martin KA, Vardaxis P, Reinemeyer C, Vijayapalani P, Robertson AP, et al. Repertoire of P-glycoprotein drug transporters in the zoonotic nematode Toxocara canis. Sci Rep. 2023;13: 4971. doi:10.1038/s41598-023-31556-1

25. James CE, Davey MW. Increased expression of ABC transport proteins is associated with ivermectin resistance in the model nematode Caenorhabditis elegans. International Journal for Parasitology. 2009;39: 213–220. doi:10.1016/j.ijpara.2008.06.009

26. Janssen IJI, Krücken J, Demeler J, von Samson-Himmelstjerna G. Caenorhabditis elegans: Modest increase of susceptibility to ivermectin in individual P-glycoprotein loss-of-function strains. Experimental Parasitology. 2013;134: 171–177. doi:10.1016/j.exppara.2013.03.005

27. Ménez C, Alberich M, Kansoh D, Blanchard A, Lespine A. Acquired Tolerance to Ivermectin and Moxidectin after Drug Selection Pressure in the Nematode Caenorhabditis elegans. Antimicrobial Agents and Chemotherapy. 2016;60: 4809–4819. doi:10.1128/aac.00713-16

28. Dube F, Hinas A, Delhomme N, Åbrink M, Svärd S, Tydén E. Transcriptomics of ivermectin response in Caenorhabditis elegans: Integrating abamectin quantitative trait loci and comparison to the Ivermectin-exposed DA1316 strain. PLOS ONE. 2023;18: e0285262. doi:10.1371/journal.pone.0285262

29. Freeman AS, Nghiem C, Li J, Ashton FT, Guerrero J, Shoop WL, et al. Amphidial structure of ivermectin-resistant and susceptible laboratory and field strains of Haemonchus contortus. Veterinary Parasitology. 2003;110: 217–226. doi:10.1016/S0304-4017(02)00321-7

30. Urdaneta-Marquez L, Bae SH, Janukavicius P, Beech R, Dent J, Prichard R. A dyf-7 haplotype causes sensory neuron defects and is associated with macrocyclic lactone resistance worldwide in the nematode parasite Haemonchus contortus. International Journal for Parasitology. 2014;44: 1063–1071. doi:10.1016/j.ijpara.2014.08.005

31. Ménez C, Alberich M, Courtot E, Guegnard F, Blanchard A, Aguilaniu H, et al. The transcription factor NHR-8: A new target to increase ivermectin efficacy in nematodes. PLOS Pathogens. 2019;15: e1007598. doi:10.1371/journal.ppat.1007598

32. Guerrero GA, Derisbourg MJ, Mayr FA, Wester LE, Giorda M, Dinort JE, et al. NHR-8 and P-glycoproteins uncouple xenobiotic resistance from longevity in chemosensory C. elegans mutants. Ünal E, Malhotra V, Ünal E, editors. eLife. 2021;10: e53174. doi:10.7554/eLife.53174

33. Dent JA, Smith MM, Vassilatis DK, Avery L. The genetics of ivermectin resistance in Caenorhabditis elegans. Proceedings of the National Academy of Sciences. 2000;97: 2674–2679. doi:10.1073/pnas.97.6.2674

34. Alberich M, Garcia M, Petermann J, Blancfuney C, Jouffroy S, Jacquiet P, et al. Evaluation of nematode susceptibility and resistance to anthelmintic drugs with a WMicrotracker motility assay. Sci Rep. 2025;15: 17968. doi:10.1038/s41598-025-02866-3

35. Fraser AG, Kamath RS, Zipperlen P, Martinez-Campos M, Sohrmann M, Ahringer J. Functional genomic analysis of C. elegans chromosome I by systematic RNA interference. Nature. 2000;408: 325–330. doi:10.1038/35042517

36. Hartman JH, Widmayer SJ, Bergemann CM, King DE, Morton KS, Romersi RF, et al. Xenobiotic metabolism and transport in Caenorhabditis elegans. J Toxicol Environ Health B Crit Rev. 2021;24: 51–94. doi:10.1080/10937404.2021.1884921

37. Wolstenholme AJ. Glutamate-gated chloride channels. J Biol Chem. 2012;287: 40232–40238. doi:10.1074/jbc.R112.406280

38. AlGusbi S, Krücken J, Ramünke S, von Samson-Himmelstjerna G, Demeler J. Analysis of putative inhibitors of anthelmintic resistance mechanisms in cattle gastrointestinal nematodes. International Journal for Parasitology. 2014;44: 647–658. doi:10.1016/j.ijpara.2014.04.007

39. Doyle SR, Laing R, Bartley D, Morrison A, Holroyd N, Maitland K, et al. Genomic landscape of drug response reveals mediators of anthelmintic resistance. Cell Reports. 2022;41: 111522. doi:10.1016/j.celrep.2022.111522

40. Laing R, Doyle SR, McIntyre J, Maitland K, Morrison A, Bartley DJ, et al. Transcriptomic analyses implicate neuronal plasticity and chloride homeostasis in ivermectin resistance and response to treatment in a parasitic nematode. PLOS Pathogens. 2022;18: e1010545. doi:10.1371/journal.ppat.1010545

41. Kamal M, Tokmakjian L, Knox J, Han D, Moshiri H, Magomedova L, et al. PGP-14 establishes a polar lipid permeability barrier within the C. elegans pharyngeal cuticle. Hart AC, editor. PLoS Genet. 2023;19: e1011008. doi:10.1371/journal.pgen.1011008

42. 42. Evans TC. Transformation and microinjection. WormBook: The Online Review of C elegans Biology [Internet]. WormBook; 2006. Available: https://www.ncbi.nlm.nih.gov/books/NBK19648/

43. Ly K, Reid SJ, Snell RG. Rapid RNA analysis of individual Caenorhabditis elegans. MethodsX. 2015;2: 59–63. doi:10.1016/j.mex.2015.02.002

44. Zhao Z, Sheps JA, Ling V, Fang LL, Baillie DL. Expression Analysis of ABC Transporters Reveals Differential Functions of Tandemly Duplicated Genes in Caenorhabditis elegans. Journal of Molecular Biology. 2004;344: 409–417. doi:10.1016/j.jmb.2004.09.052

45. Taylor SR, Santpere G, Weinreb A, Barrett A, Reilly MB, Xu C, et al. Molecular topography of an entire nervous system. Cell. 2021;184: 4329–4347.e23. doi:10.1016/j.cell.2021.06.023

46. Ghaddar A, Armingol E, Huynh C, Gevirtzman L, Lewis NE, Waterston R, et al. Whole- body gene expression atlas of an adult metazoan. Sci Adv. 2023;9: eadg0506. doi:10.1126/sciadv.adg0506

47. Cao J, Packer JS, Ramani V, Cusanovich DA, Huynh C, Daza R, et al. Comprehensive single-cell transcriptional profiling of a multicellular organism. Science. 2017;357: 661– 667. doi:10.1126/science.aam8940

48. 48. Gerhard AP. The role of nematode P-glycoproteins in the mechanisms of macrocyclic lactone resistance. Freien Universität Berlin. 2021. Available: https://refubium.fu-berlin.de/handle/fub188/31742

